# Rapid and efficient generation of human 8-cell-like cells for embryo modelling

**DOI:** 10.64898/2026.08.18.745473

**Authors:** Arda Odabas, Sıla Unlu, Eren Ozturk, Nilya Karasurmeli, Kelly Hu, Marion Leleu, Can Aztekin, Tamer T. Önder

## Abstract

8-cell blastomeres of human embryos possess broad lineage potential and undergo major zygotic genome activation (ZGA), yet experimental access to this transient cell state remains limited. Rare 8-cell-like cells (8CLCs) arise spontaneously in naïve pluripotent stem cell cultures, but their low abundance has constrained mechanistic and functional studies. Here, we develop a chemically defined strategy for rapid and robust induction of 8CLCs. Through sequential small-molecule screens focused on chromatin regulators, we identify five compounds acting through distinct pathways that generate up to 40% 8CLCs within 48 hours. The resulting cells, which we term rapidly induced 8CLCs (ri8CLCs), recapitulate key molecular features of 8-cell blastomeres, including induction of ZGA-associated genes, cleavage-stage transposable elements, and 8-cell-stage transcriptional signatures in bulk and single-cell transcriptomic analyses. Functionally, ri8CLCs exhibit enhanced developmental competence, acquiring the ability for spontaneous extraembryonic differentiation and assembly into well-cavitated blastoids on an accelerated 72-hour timeline. Notably, ri8CLC induction enables blastoid formation even in the absence of MEK inhibition, TGF-β/Activin/Nodal inhibition and exogenous LIF, revealing a developmental competence consistent with an early embryonic state. Together, these findings establish a rapid, defined, and highly efficient platform for generating human ri8CLCs and provide a tractable model for studying early human embryogenesis.

## Introduction

The 8-cell stage of human development represents a brief but pivotal period in early embryogenesis. At this stage, blastomeres are developmentally distinct from both the fertilized egg and the later lineage-restricted cells of the blastocyst^1^. They are thought to represent the last totipotent cells of the early human embryo, retaining the capacity to contribute to both embryonic and extraembryonic lineages before the first lineage decisions become consolidated. 8-cell blastomeres undergo a major wave of zygotic genome activation (ZGA), during which developmental control shifts from maternally deposited transcripts to the newly activated embryonic genome. In humans, this transition is accompanied by activation of cleavage-stage transcription factors, transposable elements, and regulatory programs that precede lineage specification^2–4^. Despite its central importance, the human 8-cell state is exceptionally transient and experimentally inaccessible, making it difficult to define how this cellular identity is established, maintained, and exited.

Rare cells resembling human 8-cell blastomeres have been identified within naïve human pluripotent stem cell (PSC) cultures and are commonly referred to as 8-cell-like cells (8CLCs)^5^. These cells typically comprise only 0.5–1% of naïve PSC cultures and are characterized by activation of 8-cell/ZGA-associated genes, including *TPRX1* and *LEUTX*, as well as cleavage-stage transposable elements^5–7^. Their spontaneous emergence suggests that naïve PSCs retain latent access to an early embryonic, totipotent-like program. However, this state appears to be unstable and short-lived in culture, and as a result, the low abundance and limited persistence of 8CLCs have constrained their use for mechanistic studies and for functional assays in embryo models.

Several studies have shown that 8-cell-like transcriptional programs can be induced *in vitro.* Genetic perturbations such as DUX4 overexpression activate cleavage-stage gene networks while chemical approaches, including inhibition of histone deacetylases and methyltransferases with trichostatin A and DZNep, increase 8CLC abundance by altering the chromatin landscape^8–10^. These observations have shown that entry into an 8-cell-like state is regulated by chromatin and transcriptional barriers. However, such approaches often involve broad epigenetic disruption and prolonged treatment, making it difficult to distinguish direct mechanisms of 8CLC induction from secondary effects of toxicity or global chromatin perturbation. A key unresolved challenge is whether the human 8-cell-like state can be induced rapidly and efficiently from human PSCs.

Here, we used human naïve PSC lines carrying knock-in TPRX1 reporters together with a curated small-molecule library to identify regulators of 8CLC induction. Because chromatin-associated processes are central to early embryonic cell-state transitions^11,12^, we focused primarily on compounds targeting epigenetic regulators. Through iterative chemical screens, we identified synergistically acting molecules and developed a multi-component cocktail capable of inducing robust 8CLC conversion, reaching up to 40% TPRX1-positive cells within 48 hours. Transcriptomic and epigenomic analyses demonstrated acquisition of an 8-cell-like identity, including activation of canonical cleavage-stage genes, increased chromatin accessibility at 8-cell regulatory elements, and induction of ZGA-associated transposable elements.

Together, our findings show that the human 8-cell-like state can be rapidly induced and identify chemical regulators of this transient state associated with expanded developmental competence. This scalable platform enables mechanistic interrogation of early human embryonic transcriptional programs and the application of 8CLCs in embryo modelling.

## Results

### Identifying inducers of 8CLC induction

To quantitatively monitor the transition from naïve pluripotency to an 8-cell-like cell (8CLC) state, we generated a TPRX1–mCherry knock-in reporter in a human induced pluripotent stem cell (iPSC) line and assessed reporter activity across pluripotent states (Fig. S1A, B). TPRX1–mCherry-positive cells were not detected under primed conditions, whereas conversion to naïve pluripotency led to the emergence of a rare reporter-positive population (∼0.9%) which could be further boosted by DUX4 overexpression, consistent with previous reports (Fig. S1C, D). Quantitative RT-PCR analysis confirmed induction of established 8CLC markers, including *DUXB*, *LEUTX*, *TPRX1*, *KLF17*, *ZNF280A*, *MBD3L2*, *ZSCAN5B* and *RFPL2*, with further enrichment of these markers in sorted mCherry-positive cells (Fig. S1E). These results validate the TPRX1–mCherry reporter as a quantitative readout of 8CLC-like transcriptional activation.

We next used this reporter to perform a focused chemical screen in naïve human PSCs (Fig. 1A). A curated library of 94 compounds targeting epigenetic and chromatin-associated pathways was screened individually by flow cytometry after 48 hours of treatment. This screen recovered several positive-control and expected activities, including histone deacetylase (HDAC) inhibitors, the HSP90 inhibitor Geldanamycin and the p53 activator Nutlin-3a, consistent with prior reports linking chromatin relaxation^7^, P-body regulation of ZGA-like transcriptional programs^13^ and stress-response pathways^6,14,15^. Nutlin-3a induced nuclear p53 accumulation, increased the proportion of TPRX1-positive cells, and activated canonical 8CLC markers in wild-type cells, whereas TP53 knockout hiPSC clones lacked p53 accumulation, were resistant to Nutlin-3a-induced cell death, and showed reporter activation and marker-gene induction only at baseline levels (Fig. S2A-D). These findings demonstrate that Nutlin-3a induces 8CLC-like states through a p53-dependent mechanism. In addition to these expected activities, several compounds targeting chromatin-associated regulatory complexes such as MEN1–MLL and LSD1/KDM1A, increased the reporter-positive population and induced the expression of canonical 8CLC markers (Fig 1B-E). Partial inhibition of splicing by Pladienolide B also increased TPRX1–mCherry activity at lower doses, although this effect was reduced at higher concentrations^16^ (Fig. 1B).

**Figure 1.**
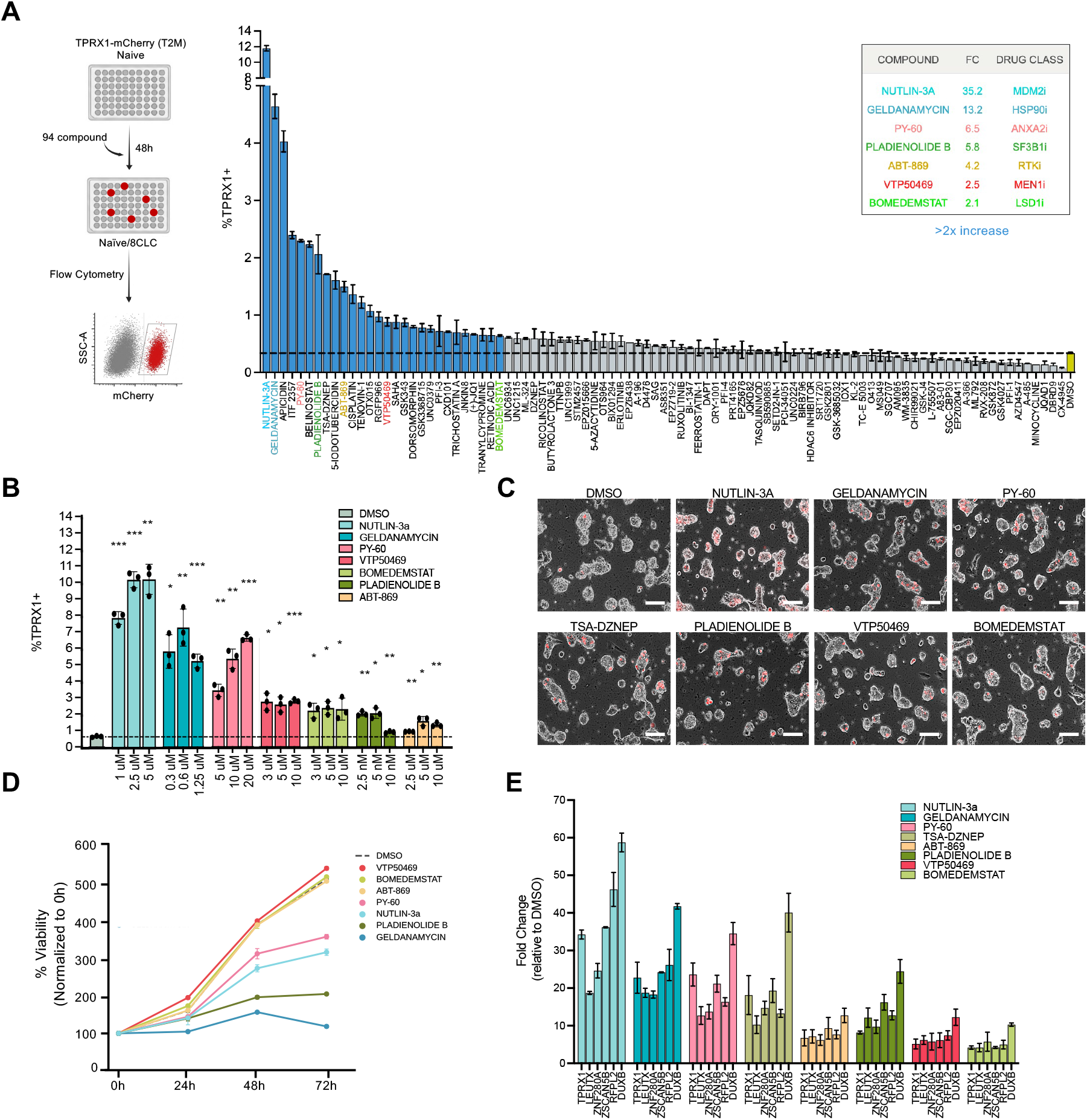
Small-molecule screening identifies regulators of 8CLC induction in naïve human pluripotent stem cells. **A**, Schematic overview of the small-molecule screening strategy and quantification of TPRX1–mCherry-positive cells following treatment with individual compounds from the primary screen. Selected compounds associated with increased reporter activation are highlighted. Dashed line indicates the DMSO threshold used to define hit compounds. The inset table summarizes representative candidate compounds and their annotated target classes**. B,** Dose-response validation of selected primary-screen hits. Naive TPRX1-mCherry reporter cells were treated for 48 h with vehicle (DMSO) or with the indicated compounds at three concentrations each. Dashed line indicates the mean of DMSO-treated controls**. C,** Representative images of naïve TPRX1–mCherry reporter cells following treatment with indicated compounds. Scale bars, 200 μm. **D,** Cell viability analysis of naïve cells treated with selected compounds over a three-day time course, relative to DMSO-treated controls. **E,** RT–qPCR analysis of 8CLC-associated marker genes following treatment with selected compounds relative to DMSO-treated controls.

Among the non-HDAC candidates, PY60, a compound previously reported to act as a YAP agonist^17^, emerged as a particularly robust inducer of TPRX1-positive cells, increasing their prevalence in a dose-dependent manner up to 7% (Fig. 1B). PY60 induced multiple 8CLC-associated genes, including *TPRX1*, *LEUTX*, *ZNF280A*, *ZSCAN5B*, *RFPL2* and *DUXB*, while maintaining comparatively higher cell viability than Nutlin-3a, Geldanamycin or Pladienolide B over a three-day treatment period (Fig. 1D, E). We next asked whether PY60-mediated 8CLC induction depends on the pluripotent state or on a specific naïve culture condition. PY60 failed to induce detectable TPRX1–mCherry expression in primed PSCs, indicating that its activity requires a naïve state. However, within naïve PSCs, PY60 induced 8CLCs across multiple culture conditions, including HENSM^18^, PXGL^19^ and 5iLA^20^ (Fig. S2E, F). Together, these findings identify PY60 as a potent and relatively well-tolerated inducer of 8CLC conversion that acts within a naïve pluripotent context but is not dependent on a specific naïve culture condition.

### PY60 induces an 8CLC transcriptional program

To further characterize the transcriptional changes induced by PY60, we performed RNA sequencing following 24-hour and 48-hour treatment, as well as in TPRX1-positive cells isolated after 48 hours (Fig. 2A). As a positive control, DUX4-overexpressing cells were sorted and analyzed in parallel. We compared PY60 induced transcriptional profiles to stage-specific gene signatures derived from human preimplantation embryos. PY60-treated cells showed progressive activation of genes associated with the 4-cell and 8-cell stages, with increasing enrichment from 24 hours to 48 hours and further in the sorted mCherry-positive population, indicating a gradual acquisition of a ZGA-like transcriptional state (Fig. 2A). Differential expression analysis confirmed strong upregulation of key 8-cell-associated genes, including *LEUTX*, *DUXB*, *ZSCAN* family members and *MBD3L2*, in both PY60-treated and DUX4-overexpressing cells with comparable magnitudes of induction (Fig. 2B). Gene set enrichment analysis (GSEA) using an 8-cell-stage signature^5^ demonstrated significant enrichment in PY60-treated cells compared to DMSO controls, similar to DUX4 overexpression (Fig. 2C). GSEA across the complete MSigDB collection identified the 8C–Morula signature as the most strongly enriched program in PY60-sorted cells, exhibiting the highest normalized enrichment score among all significantly enriched gene sets (Fig. 2D). Conversely, genes associated with pluripotency were significantly depleted in both PY60-treated and DUX4-overexpressing cells, indicating a transcriptional shift away from the pluripotent state and toward an 8CLC identity (Fig. 2E). Canonical DUX4 target genes, including *DUXB*, *LEUTX*, PRAMEF family members and *ZSCAN* genes were highly induced in PY60-treated cells (Fig. 2F). In addition, PY60 treatment induced expression of multiple transposable elements associated with human ZGA and 8-cell-stage embryos, including MLT2A1, MLT2A2, and HERV/LTR-family elements^21,22^ (Fig. 2G). These findings further support activation of an 8-cell-like transcriptional state following PY60 treatment.

**Figure 2.**
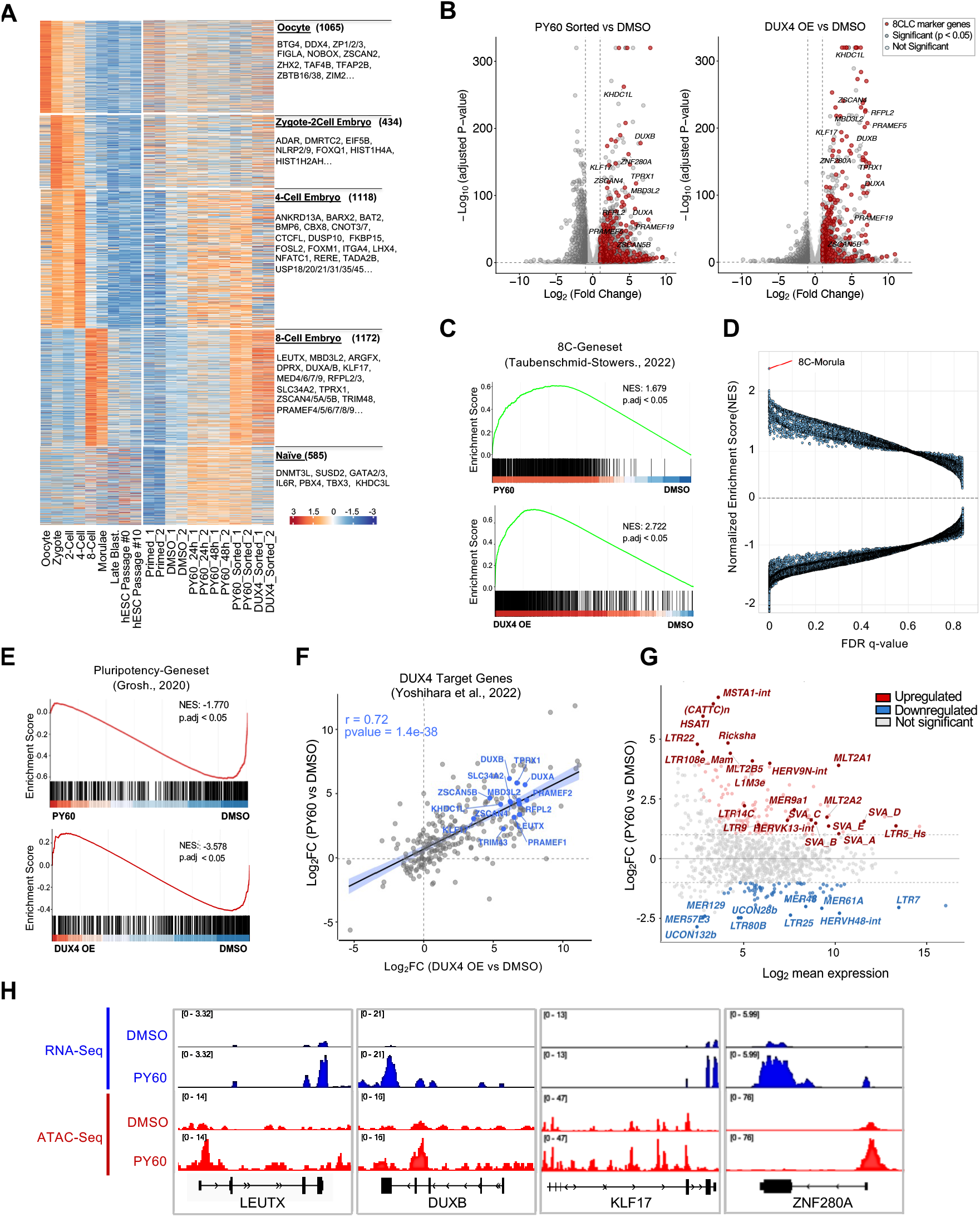
PY60 induces a ZGA-like transcriptional program. **A**, Stage-specific gene expression signatures across human preimplantation developmental stages and experimental conditions. **B,** Differential gene expression in PY60-sorted and DUX4-overexpressing cells relative to DMSO-treated controls. Representative 8CLC-associated genes are highlighted. **C,** Gene set enrichment analysis (GSEA) using a published 8-cell-stage gene signature in PY60-treated and DUX4-overexpressing cells relative to DMSO controls. NES, normalized enrichment score. **D**, GSEA performed against all gene sets in the MSigDB database. Each dot represents a gene set. **E,** GSEA using a pluripotency gene signature (Grosh et al., 2020). **F,** Log₂ fold changes in DUX4 target genes (Yoshihara et al., 2022) between PY60-sorted versus DMSO and DUX4-overexpressing versus DMSO conditions. Pearson correlation coefficient is shown in blue. Representative ZGA-associated genes are highlighted. **G,** MA plot showing differential expression of transposable element families following PY60 treatment relative to DMSO controls. Upregulated elements are shown in red; downregulated elements are shown in blue. Representative transposable element families associated with human ZGA, including MLT2A1, MLT2A2 and HERVH-family elements, are labelled. **H,** Genome browser tracks showing ATAC-seq and RNA-seq signal at representative ZGA-associated loci (*LEUTX, DUXB, KLF17,* and *ZNF280A*) in DMSO-and PY60-treated cells.

We next asked whether PY60-induced transcriptional changes were accompanied by remodeling of chromatin accessibility. Assay for Transposase-Accessible Chromatin using sequencing (ATAC-seq) revealed widespread changes in accessible chromatin following PY60 treatment, including increased accessibility at regulatory regions associated with canonical 8CLC genes such as *LEUTX*, *DUXB* and *ZNF280A* (Fig. 2H and Fig. S3A). Motif enrichment analysis of gained regions identified strong enrichment for CTCF, CTCFL (BORIS), TFAP2 and NFE2L2 motifs, whereas regions losing accessibility were enriched for pluripotency-associated transcription factor motifs such as OCT4 and SOX2. Notably, approximately 64% of regions gaining accessibility overlapped transposable element loci, suggesting that PY60-mediated induction of 8CLC states may involve removal of transcriptional repression at TE-associated genomic regions linked to human ZGA programs (Fig. S3B). Collectively, these findings indicate that PY60 induces not only an 8-cell transcriptional program, but also a chromatin accessibility landscape enriched at cleavage-stage and transposable element-associated regulatory regions.

PY60 was previously identified as a molecule that promotes YAP activation through disruption of the ANXA2–LATS1/2 interaction^17^. We therefore sought to determine whether the transcriptional effects observed during 8CLC induction were linked to canonical ANXA2/YAP signaling. Gene set enrichment analysis revealed modest enrichment of genes bound by YAP^23^ following PY60 treatment; however, enrichment of established YAP target genes^24^ was not statistically significant (Fig. S3C), suggesting that PY60-induced 8CLC conversion may not be primarily driven by canonical YAP transcriptional activity. To determine whether PY60-mediated 8CLC induction depends on canonical ANXA2–YAP signaling, we depleted either ANXA2 or YAP1 using CRISPR-mediated gene targeting. Loss of ANXA2 did not significantly affect the proportion of TPRX1-positive cells either at baseline or following PY60 treatment compared with non-targeting controls (Fig. S3D, E). Similarly, YAP1 knockout neither phenocopied PY60 treatment nor attenuated PY60-mediated induction of TPRX1-positive cells (Fig. S3F, G). To test whether alternative means of activating YAP could recapitulate the PY60 phenotype, we treated naïve TPRX1 reporter cell line with the LATS1/2 inhibitors, TRULI and GA-017. In contrast to PY60, both compounds reduced the proportion of TPRX1-positive cells and induced morphological differentiation (Fig. S3H, I). These findings are consistent with previous reports showing that excessive YAP activation promotes trophectoderm-like differentiation in naïve pluripotent stem cells^25,26^ and further support the conclusion that PY60-induced 8CLC conversion is mechanistically distinct from canonical Hippo/YAP activation.

### Rapid and robust chemical induction of 8CLCs

To determine whether PY60-induced 8CLC conversion could be further enhanced, we performed a secondary screen to identify compounds that can act in concert with PY60 (Fig. 3A). Naïve TPRX1–mCherry reporter cells were treated with individual compounds in combination with PY60, and the proportion of mCherry-positive cells was quantified by flow cytometry. Among the top hits, 5-azacytidine, a DNA methyltransferase inhibitor, showed a strong effect in combination with PY60. However, even at low concentrations, 5-azacytidine markedly reduced cell viability, limiting its utility for generating 8CLC populations (Fig. S4A). Several compounds previously implicated in inducing totipotent-like cells such as DOT1L inhibitor EPZ-5676^27,28^, METTL3 inhibitor STM2457^29^, SUMOylation inhibitor ML-792^30^ did not have additive effects with PY60. However, Retinoic acid (RA) and TTNPB, an RA analog, showed a clear effect when combined with PY60, almost doubling the fraction of mCherry-positive cells (Fig. 3A). We confirmed that this effect was accompanied by increased expression of 8CLC marker genes, indicating that the observed increase reflects activation of a ZGA-like transcriptional program rather than non-specific reporter-activation (Fig. S4B). In addition, we followed up on compounds not previously linked to 8CLC state, including JQKD82, a KDM5 inhibitor and Bomedemstat, a KDM1A inhibitor. To define the final chemical induction recipe, we performed omission experiments where all candidate compounds identified from both screens were applied in combination, followed by sequential removal of individual components. The complete combination produced the highest frequency of TPRX1-positive cells, whereas omission of PY60 or retinoic acid caused the most pronounced reduction in 8CLC induction, and individual removal of Nutlin-3a, Bomedemstat, or JQKD82 led to more modest decreases (Fig. 3B).

**Figure 3.**
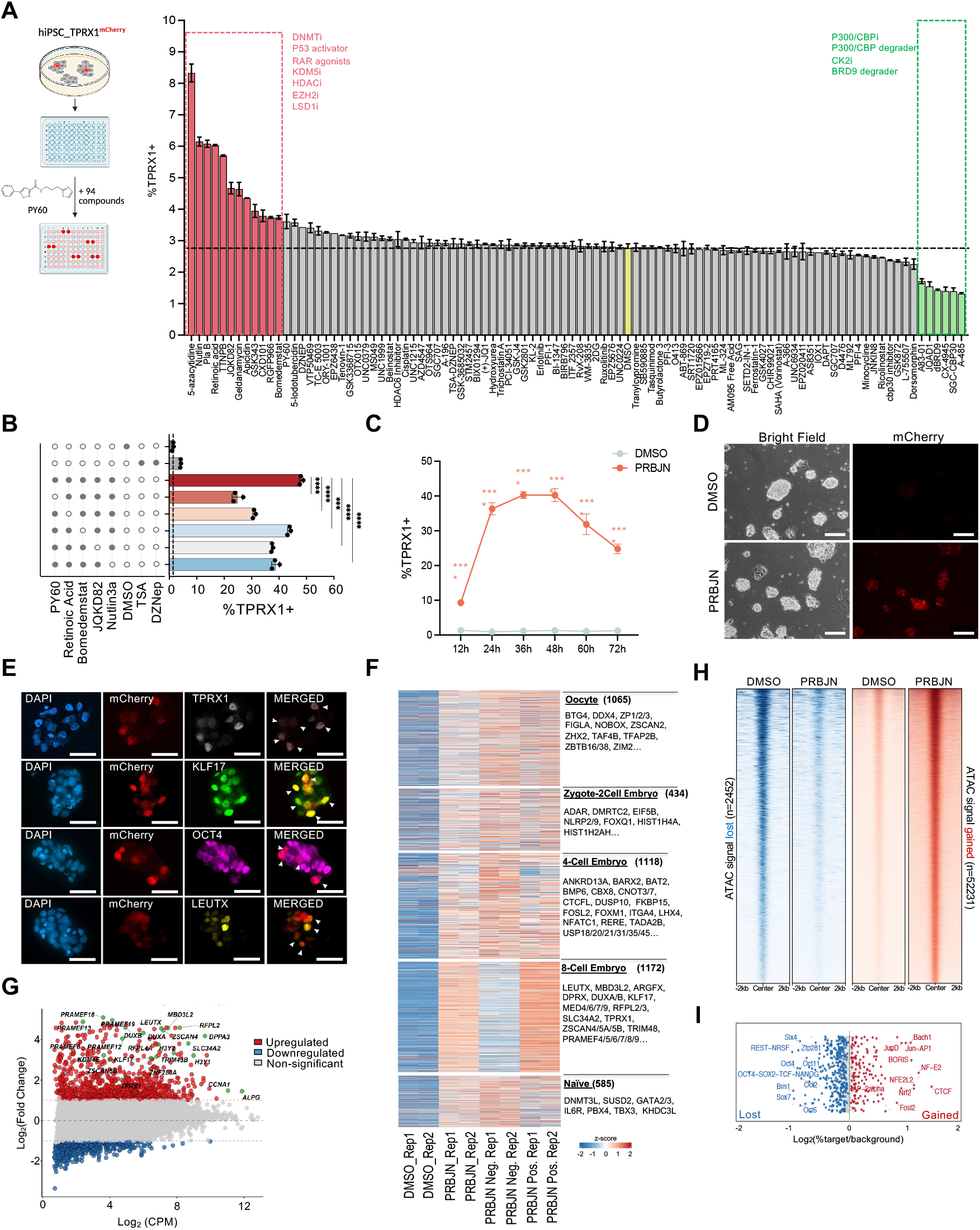
Defined chemical cocktail enhances PY60-driven induction of 8CLC-like cells. **A**, Overview of the secondary chemical screen and flow cytometry results. Hit compounds that increase (red) or decrease PY60 effect (green) are highlighted. The dashed line and the yellow bar indicate PY60 alone. **B,** % TPRX1-poistive cells upon omission of individual components from the full compound combination. Filled circles indicate compounds present in each condition. **p < 0.01, ***p < 0.001, ****p < 0.0001, one-way ANOVA;. **C,** Time-course analysis of TPRX1–mCherry induction following treatment with DMSO or the PRBJN cocktail. **p < 0.01, ***p < 0.001, ****p < 0.0001, two-way ANOVA. **D,** Representative bright-field and mCherry fluorescence images of naïve TPRX1–mCherry reporter cells treated with DMSO or PRBJN for 24 hours. Scale bars, 200 μm. **E,** Immunofluorescence staining of PRBJN-treated cells for 8CLC-associated markers, KLF17, LEUTX, TPRX1, and pluripotency marker OCT4. Scale bars, 50 μm. **F,** Stage specific genes in DMSO/PRBJN-treated bulk populations and sorted mCherry-positive and negative populations. **G,** MA plot showing differential gene expression following PRBJN treatment relative to DMSO. **H,** Heatmaps of ATAC-seq signal centered on differentially accessible regions in DMSO-versus PRBJN-treated cells, ranked by signal intensity and displayed within ±2 kb of the peak center. Left (blue): regions that lose accessibility upon PRBJN treatment (ATAC signal lost, n = 2,452). Right (red): regions that gain accessibility (ATAC signal gained, n = 52,231). **I,** Transcription factor motif enrichment for the lost (blue) and gained (red) regions, plotted as log₂(% target/background)

Based on these results, we defined a five-component cocktail (PRBJN) and assessed its ability to induce 8CLCs over time. The final PRBJN cocktail consisted of <u>P</u>Y60 (7.5 μM), <u>R</u>etinoic acid (0.5 μM), <u>B</u>omedemstat (1 μM), <u>J</u>QKD82 (1 μM) and <u>N</u>utlin-3a (0.5 μM). Treatment with PRBJN resulted in rapid induction of TPRX1-positive cells, reaching ∼40% within 48 hours (Fig. 3C, D). In contrast, treatment of naïve PSCs with TSA (20 nM), DZNEP (50 nM) for the same duration resulted in 6% TPRX1-positive cells (Fig. 3B). PRBJN treatment induced robust reporter activation as well as TPRX1 protein expression as assessed by immunofluorescence staining (Fig 3E). We also observed that mCherry-positive 8CLCs displayed reduced OCT4 but higher LEUTX and KLF17 expression (Fig. 3E). Importantly, similar levels of 8CLC induction were observed in additional human pluripotent stem cell lines, including H9 and WIBR3, indicating that the 8CLC-inducing activity of PRBJN is reproducible across multiple genetic backgrounds and is not restricted to iPSCs or ESCs (Fig. S4C). Collectively, these results demonstrate that PRBJN enables rapid, robust, and broadly applicable induction of the 8CLC state, substantially outperforming previously reported chemical induction approaches.

To determine whether PRBJN treatment induced transcriptional features associated with human 8-cell blastomeres, we performed RNA sequencing on PRBJN-treated bulk populations, as well as sorted mCherry-positive and mCherry-negative fractions (Fig. 3F). Stage-specific gene signatures showed progressive enrichment of 8-cell-stage transcriptional program, particularly within the sorted mCherry-positive population (Fig. 3F). Consistent with this, differential expression analysis revealed strong upregulation of canonical 8CLC-associated genes, including DUX-family targets, PRAMEF family members, *LEUTX*, *ZSCAN* genes and *DPPA3*, confirming activation of a ZGA-like transcriptional program following PRBJN treatment (Fig. 3G).

We next investigated whether PRBJN-induced transcriptional changes were accompanied by chromatin remodeling. ATAC-seq analysis revealed extensive changes in chromatin accessibility following PRBJN treatment, with 52,231 regions gaining accessibility and 2,452 regions losing accessibility relative to DMSO controls (Fig. 3H). Regions gaining accessibility were enriched for motifs associated with AP-1 family members, CTCF/CTCFL, NF-E2 and BACH1, whereas regions losing accessibility were enriched for pluripotency-associated motifs, including OCT4/SOX2/TCF/NANOG and SOX family motifs (Fig. 3I). Together, these data indicate that PRBJN rapidly induces an 8CLC-like transcriptional and chromatin state while reducing features associated with naïve pluripotency.

To further determine whether PRBJN-induced 8CLCs transcriptionally resemble human 8-cell-stage blastomeres, we performed single-cell RNA sequencing of naïve TPRX1– mCherry cells treated with either DMSO or PRBJN for 24 hours. Unsupervised clustering identified nine transcriptionally distinct cell populations, including a cluster markedly enriched following PRBJN treatment (cluster 4), while smaller numbers of transcriptionally similar cells were also detected in DMSO-treated samples, consistent with endogenous 8CLC populations previously observed in naïve pluripotent cultures^5–7^ (Fig. 4A). Cluster 4 displayed strong enrichment of canonical 8CLC-associated genes, including *TPRX1*, *LEUTX*, *DUXA*, *H3Y*, *ZSCAN4*, *KHDC1L*, *DPPA3*, *SLC34A2* and multiple PRAMEF family members (Fig. 4B-C). In contrast, core pluripotency-associated genes such as *POU5F1* and *SOX2* were reduced (Fig. 4D).

**Figure 4.**
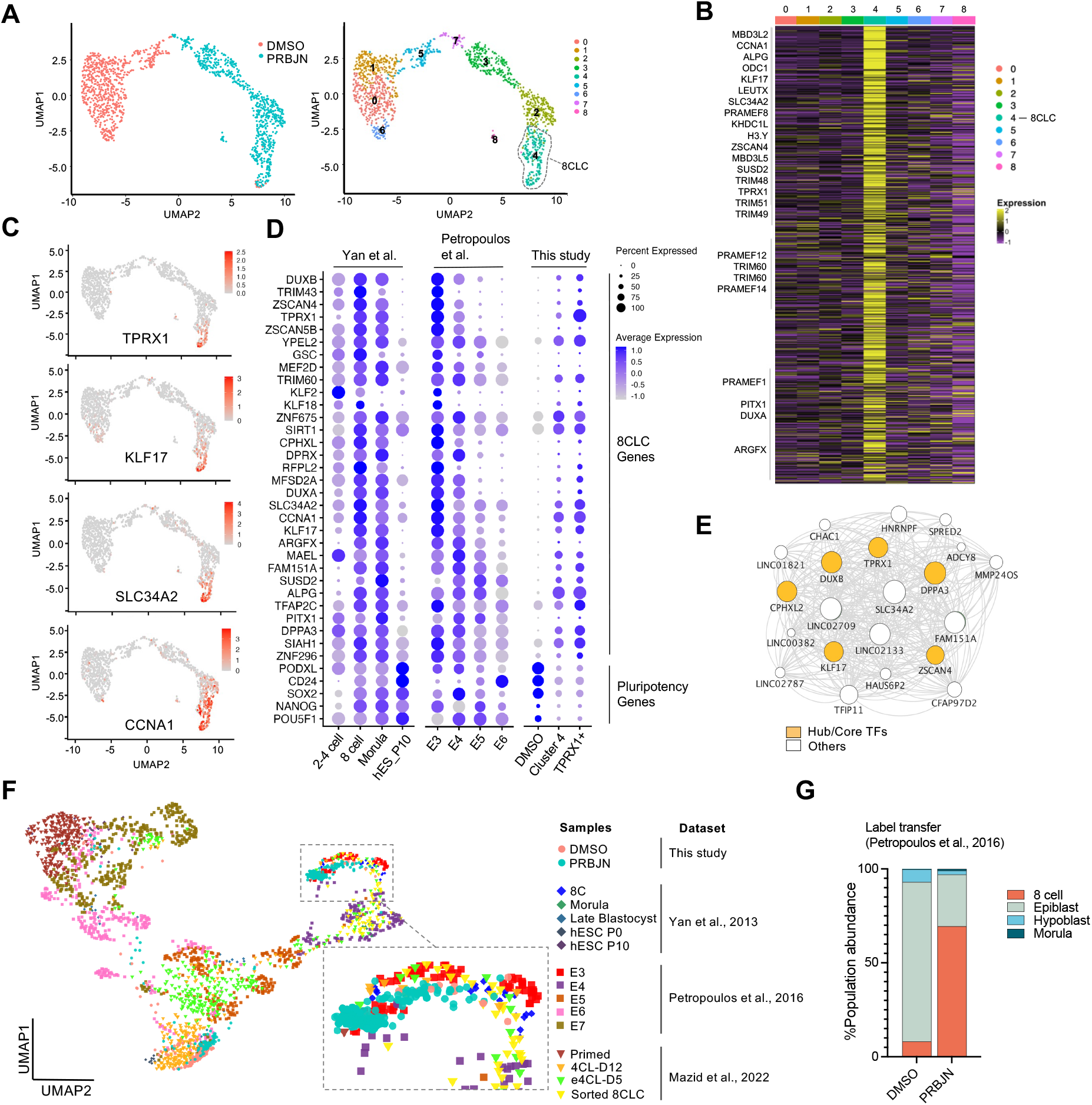
Single-cell transcriptomic analysis reveals emergence of an 8CLC-like population upon PRBJN treatment. **A**, UMAP representation of naïve TPRX1–mCherry reporter cells treated with DMSO or PRBJN for 24 h, colored by treatment group (left) or unsupervised cluster identity (right). **B,** Heatmap showing scaled expression of representative marker genes across identified clusters. Cluster 4 displays strong enrichment of canonical 8CLC/ZGA-associated genes, including *TPRX1, LEUTX, ZSCAN4, KHDC1L, CCNA1, DUXA/B and PRAMEF* family members. **C,** Feature plots showing expression of representative 8CLC markers (*TPRX1, KLF17, SLC34A2,* and *CCNA1*) across the dataset. **D,** Comparative expression analysis of representative 8CLC-associated genes across published human embryo datasets (Yan et al., 2013; Petropoulos et al., 2016) and cells from this study. Dot size indicates percentage of expressing cells and dot color indicates average expression level. **E,** Gene regulatory network analysis identifying a co-expression module enriched for canonical 8CLC-associated genes in PRBJN-induced cells. **F,** Integrated UMAP projection of the current scRNA-seq dataset with published human embryo datasets (Yan et al., 2013; Petropoulos et al., 2016; Mazid et al., 2022). Cells are colored by sample and dataset of origin as indicated. **G,** Stacked bar chart showing the predicted developmental stage assignments of DMSO and PRBJN-treated cells following label transfer from the Petropoulos et al., 2016 human embryo dataset. The y-axis represents the percentage of cells predicted to belong to each embryonic cell type.

To assess the developmental identity of these cells, we compared expression profiles from our dataset with published human embryo single-cell RNA-seq datasets from Yan et al., 2013^31^ and Petropoulos et al., 2016^32^ (Fig. 4D). Cluster 4 and TPRX1-positive cells showed transcriptional profiles closely resembling human 8-cell-stage embryos, including expression of canonical ZGA-associated genes. Gene regulatory network analysis^33^ further identified a coordinated co-expression module enriched for canonical 8CLC-associated genes, including *TPRX1*, *DUXB*, *DPPA3*, *KLF17* and *ZSCAN4*, indicating activation of an organized ZGA-like transcriptional network within the PRBJN-induced population (Fig. 4E).

Finally, to directly compare our cells with previously published embryonic and 8CLC datasets, we integrated our scRNA-seq data with human embryo datasets and previously reported 8CLC datasets^10,31,32^. Reference mapping revealed that PRBJN-treated cells localized to regions occupied by human 8-cell-stage embryos, whereas DMSO-treated cells contributed to this region at substantially lower frequencies (Fig. 4F). Label-transfer analysis assigned PRBJN-induced cells but not DMSO controls, predominantly to an 8-cell-like identity, based on Petropoulos et al. dataset (Fig. 4G). Together, these analyses demonstrate that PRBJN treatment robustly and rapidly induces cells transcriptionally resembling human 8-cell-stage embryos within 24 hours and therefore, we term these cells rapidly induced 8CLCs, or ri8CLCs.

### ri8CLCs exhibit enhanced developmental potential

To determine whether ri8CLCs have increased competence for extraembryonic lineage differentiation, we assessed their ability to undergo trophoblast lineage differentiation without external stimuli (Fig. 5A). A GATA3–mKO2 reporter naïve iPSC line was treated with either DMSO or PRBJN for 24 hours and subsequently transferred to N2B27 basal medium for an additional four days to allow for spontaneous differentiation in the absence of strong trophectoderm inducing signals such as MEK and Nodal inhibitors^34^ (Fig. 5A). Compared with DMSO controls, PRBJN pretreatment resulted in a marked increase in the proportion of GATA3-positive cells in N2B27 medium after 96 hours (Fig. 5B, S5A). While spontaneous differentiation of DMSO-treated cells generated only a small GATA3-positive population (∼9%), PRBJN-pretreated cells produced a substantially larger GATA3-positive fraction (∼40%), representing more than a four-fold increase in trophoblast-like differentiation efficiency under minimal conditions. Consistent with these findings, qPCR analysis showed downregulation of 8CLC markers and robust induction of trophectoderm-associated genes following differentiation of PRBJN-treated cells, with markers of trophoblast specification, including *GATA3*, *NR2F2* and *TP63*, expressed at significantly higher levels than in DMSO-treated controls (Fig. 5C, D). Upon further differentiation, PRBJN-treated cells, were able to generate syncytiotrophoblasts (STBs) and extravillous trophoblasts (EVTs), as demonstrated by the expression *GCM1*, *ENDOU*, *CGB*, *CGA*, *SDC1* in the case of STBs and *KRT7*, *ADAM12*, *HTRA4*, *MMP2*, *HLA-G* in the case of EVTs (Fig. 5E-G). In addition, PRBJN-derived TSCs were able to generate STBs that release of human chorionic gonadotropin into culture medium whereas DMSO-derived cells failed to do so (Fig. 5H). Collectively, these results demonstrate that ri8CLCs display extraembryonic differentiation capacity *in vitro*.

**Figure 5.**
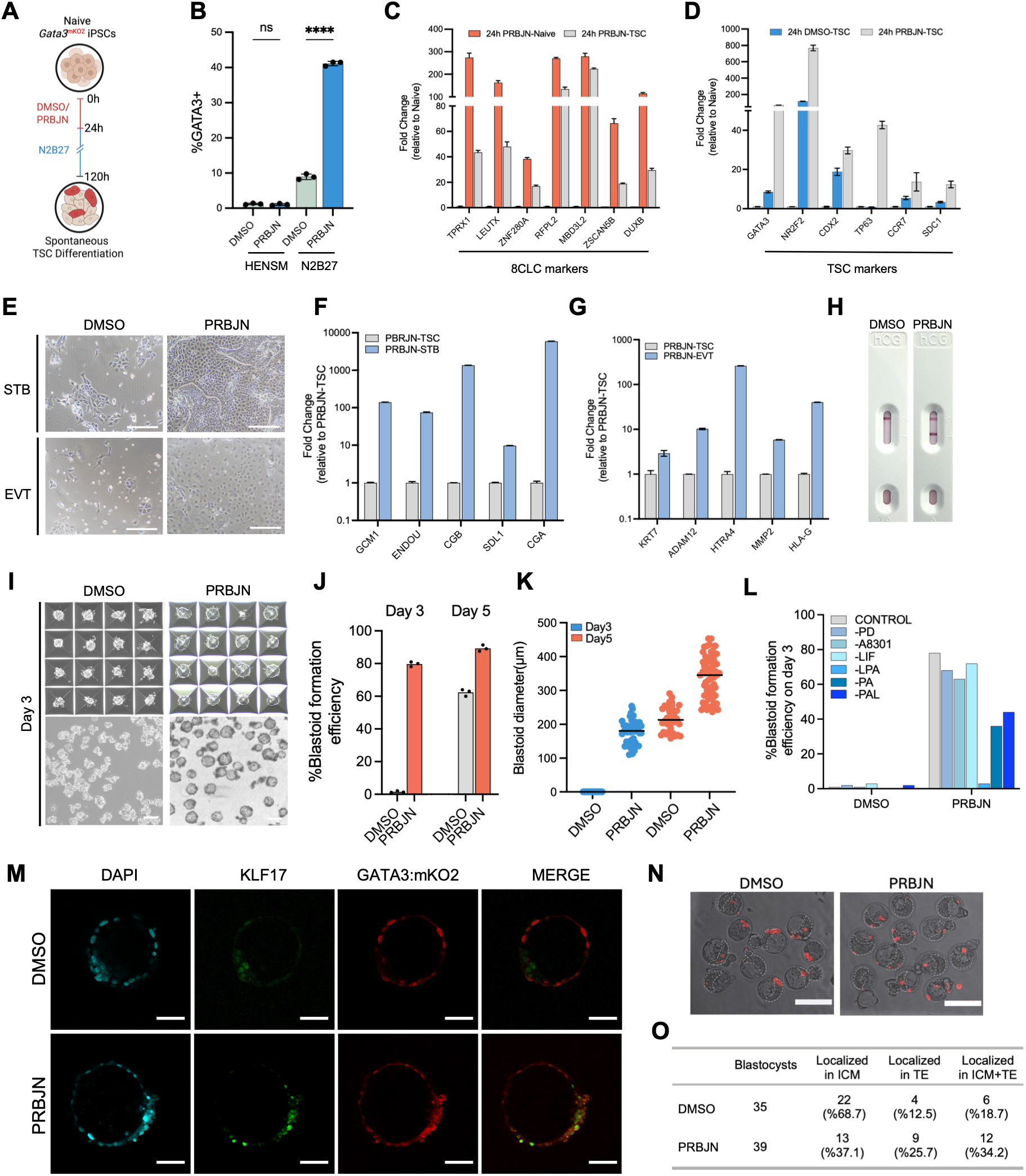
PRBJN-induced cells exhibit enhanced developmental potential. **A**, Naïve GATA3–mKO2 iPSCs were treated with DMSO or PRBJN for 24 h followed by transfer to N2B27 basal medium for four days to allow spontaneous trophoblast stem cell (TSC) differentiation. **B,** Proportion of GATA3-positive cells across conditions. ns, not significant; ***p < 0.001; ****p < 0.0001, one-way ANOVA. **C,** RT–qPCR analysis of 8CLC genes in upon 24 h PRBJN treatment and subsequent TSC differentiation **D,** RT–qPCR analysis of trophoblast stem cell genes in naïve cells, 24 h DMSO-derived TSC (DMSO-TSC), and 24 h PRBJN-derived TSC (PRBJN-TSC). **E,** TSC-derived syncytiotrophoblast (STB) and extravillous trophoblast (EVT) differentiation cultures generated from DMSO-and PRBJN-treated cells. Scale bars, 200 μm. **F,** RT-qPCR analysis of STB-specific marker genes. **G,** RT-qPCR analysis of EVT-specific marker genes following differentiation **H,** Human chorionic gonadotropin (hCG) pregnancy test results performed on conditioned media collected from differentiated cultures. **I,** Representative bright-field images of blastoids generated from naïve hPSCs following DMSO or PRBJN pre-treatment prior to aggregation. Scale bars, 200 μm. **J,** Blastoid formation efficiency on day 3 and day 5. Efficiency was calculated as the percentage of microwells containing morphologically identifiable blastoids relative to the total number of seeded aggregates. **K,** Dot plot showing the diameter (μm) of blastoids formed under DMSO and PRBJN treatment conditions at Day 3 (blue) and Day 5 (orange). Each dot represents an individual blastoid measurement. **L,** Bar chart showing the percentage blastoid formation efficiency at Day 3 under DMSO and PRBJN treatment conditions following the individual withdrawal of PALLY medium components. **M,** Representative immunofluorescence images of blastoids generated under PRBJN (top) and DMSO (bottom) conditions, stained for NANOG (green; epiblast), SOX17 (yellow; hypoblast/primitive endoderm) and GATA3-mKO2 (red; trophectoderm), with nuclei counterstained with DAPI (blue). Scale bars, 50 μm. **N,** Representative bright-field and H2B–mCherry fluorescence overlay images of mouse blastocysts following injection of DMSO-or PRBJN-pretreated human naïve PSCs. Scale bars, 100 μm. **O,** Quantification of the localization of H2B-mCherry-positive human cells within the ICM, TE, or both compartments (ICM+TE) of host mouse blastocysts. Numbers of scored blastocysts: DMSO, n = 35; PRBJN, n = 39.

We next asked whether this increased extraembryonic differentiation ability would endow ri8CLCs with a superior ability to generate early embryo models termed blastoids. Blastoid formation from human PSCs using the PALLY protocol typically takes 120 hours from initial cell seeding34. Using this protocol, PRBJN-treated cells formed well-cavitated structures by 72 hours with an efficiency approaching 80% and a mean diameter of 177 ± 34 µm, whereas DMSO-treated naïve cells remained as smaller aggregates with little or no cavitation (Fig 5I-K). As expected, DMSO-treated naïve cells formed blastoids only after 5 days of incubation. Given this rapid formation of blastoids, we next asked if components of the PALLY protocol could be omitted, yielding a minimal set of conditions that would support blastoid formation. PBRJN treatment rendered MEK inhibition (PD0325902), TGF-β/Activin/Nodal inhibition (A83-01) and exogenous LIF supplementation completely dispensable for successful blastoid formation while LPA remained necessary. Notably, PBRJN-treated cells could still form cavitated structures when all three of the dispensable components were omitted (Fig 5L, S5B). Blastoids contained inner cells expressing NANOG surrounded by GATA3-positive trophectoderm (Fig 5M).

A functional hallmark of totipotent blastomeres is their unrestricted ability to integrate into both the ICM and trophectoderm (TE) when introduced into a host embryo. To determine whether ri8CLCs possess this type of expanded developmental competence, we tested whether PRBJN pretreatment enhances the chimeric contribution of naïve PSCs toward both embryonic and extraembryonic lineages upon injection into mouse blastocysts. H2B–mCherry expressing naïve PSCs were treated with either DMSO or PRBJN for 12 hours, dissociated to single cells, and injected at a density of 8–10 cells per embryo into mouse embryos at the 8-cell stage. Injected embryos were cultured to the blastocyst stage and the localization of H2B–mCherry-positive cells within the ICM, TE, or both compartments was scored (Fig. 5N, O). Both DMSO-and PRBJN-treated cells were capable of contributing to the ICM and TE; however, PRBJN-pretreated cells showed higher overall chimerism and a substantially greater proportion of TE (12.5% vs 27.5%) and dual ICM+TE (18.7% vs 34.2%) contributions, indicating that PRBJN treatment shifts the developmental bias of human naïve PSCs toward a broader, lineage-unrestricted distribution consistent with totipotent-like competence. Together, these findings demonstrate that ri8CLCs possess a rewired lineage competence and closely mimic the developmental flexibility of 8-cell blastomeres.

## DISCUSSION

Zygotic genome activation (ZGA) represents a fundamental transition in early embryogenesis yet modelling this state in vitro using naïve PSC-derived 8-cell-like cells (8CLCs) has been limited by the rarity and transient nature of these cells. In this study, we establish a chemically defined strategy for rapid and efficient induction of human 8CLCs from naïve PSCs. Through chromatin-focused chemical screening, we identified a small-molecule combination that induces up to ∼40% TPRX1-positive cells within 48 hours. The resulting rapidly induced 8CLCs (ri8CLCs) recapitulate key molecular features of human 8-cell blastomeres, including activation of ZGA-associated genes, DUX4 targets, cleavage-stage transposable elements and 8-cell-stage transcriptional signatures in bulk and single-cell transcriptomic analyses. Functionally, ri8CLCs also display expanded developmental competence, as they efficiently generate trophoblast-like cells under minimal differentiation conditions and accelerate blastoid formation compared with naïve PSCs.

A major implication of our study is that multiple barriers prevent entry into the 8CLC state. The PRBJN cocktail combines PY60 with Nutlin-3a, retinoic acid, the KDM1A inhibitor Bomedemstat, and the KDM5 inhibitor JQKD82. We demonstrate that the effects of Nutlin-3a are p53-dependent, as p53-knockout cells fail to induce 8CLCs upon Nutlin-3a treatment. The role of retinoic acid agonists is consistent with the known activity of this pathway in inducing 2-cell-like cells (2CLCs) from mouse ESCs^36^. How Bomedemstat, JQKD82, and PY60 increase the frequency of human 8CLCs remains to be fully determined, but several mechanisms are likely involved. Bomedemstat targets KDM1A, which represses endogenous retroviruses and neighboring genes in mouse ESCs, including MERVL-associated two-cell/ZGA genes^37,38^. Additionally, KDM1A functions within the CoREST and NuRD corepressor complexes, suggesting that Bomedemstat may promote 8CLC emergence by relieving a combined demethylation/deacetylation barrier at TE-derived regulatory elements^39^. Our ATAC-seq data reinforces this notion, revealing over 52,000 regions gaining accessibility upon PRBJN treatment, with a striking 64% localization to TE loci. Concurrently, JQKD82-mediated inhibition of the KDM5 family^40^ likely elevates local H3K4 trimethylation (H3K4me3) marks at these newly accessible loci, creating a transcriptionally permissive state that supports the robust expression of 8CLC genes. Finally, the ri8CLC cocktail includes PY60, which on its own can rapidly induce an 8CLC gene signature in naïve PSCs. Although initially identified as a YAP agonist, our genetic knockout data point to a YAP-independent role in 8CLC induction, meaning the precise mechanism of PY60 remains to be explored. Taken together, these results highlight multiple, distinct pathways that actively repress the 8CLC state in naïve PSCs.

We observed that ri8CLCs are able to give rise to trophectodermal lineages *in vitro* without modulators of extracellular signals. This finding indicates that, once exiting from the 8-cell-like state, ri8CLCs can undergo spontaneous extraembryonic differentiation. This ability has not been previously observed in naïve PSCs or 8CLCs generated by alternative methods ^7,41,42^. In addition, PRBJN treatment endows naïve cells with robust and accelerated blastoid-forming capacity that closely recapitulates the timing of blastocyst formation from the embryonic 8-cell stage. Chimeric blastocyst injection experiments further supported the expanded developmental potential of PRBJN-treated cells. Whereas naïve PSCs typically contribute exclusively to the ICM in chimera assays, PRBJN-treated cells contributed to both the ICM and TE, a capacity associated with totipotent blastomeres, indicating that PRBJN treatment endows cells with a broader developmental competence consistent with a totipotent-like identity.

Together, our results define a scalable and chemically controlled platform for inducing human 8CLCs and provide insights into the barriers that restrict entry into ZGA-like states. The identification of a rapid and robust pathway for 8CLC induction, combined with the ability to functionally validate this state, establishes a framework for studying early human development and its dysregulation.

## Methods

### Cell culture

Primed human pluripotent stem cells were maintained in mTeSR medium and cultured on Geltrex-coated tissue culture plates in a 5% CO_2_ and 5% O_2_ incubator. For naïve conversion cells were dissociated with Accutase and seeded at 30% confluency with 10 μM Y-27632. Medium was refreshed the next day with HENSM^43^ (1μM PD0325901, 2 μM XAV939, 1 μM Sotrastaurin, 1 μM CGP77675, 0.8 μM BIRB0796 and 20 ng/mL human LIF in N2B27 basal medium. Naïve PSCs were maintained under hypoxic conditions and passaged every 3-4 days using 5 min incubation with Trypsin. Medium was replaced daily throughout the experiments. For media comparison experiments, cells were adapted for at least three passages in HENSM, PXGL^44^ (in which Gö6983 was replaced with an alternative PKC inhibitor, Sotrastaurin) or 5i/L/A^45^ naïve culture conditions before compound treatment. All cultures were routinely tested and confirmed negative for mycoplasma contamination.

### Generation of knock-in reporter cells

To generate a reporter system for monitoring 8-cell-like cell (8CLC) induction, a TPRX1– mCherry knock-in cell line was established using CRISPR–Cas9-mediated homologous recombination. A donor construct containing a P2A-GFP reporter cassette was kindly provided by Miguel A. Estaban^7^. GFP cassette was replaced with mCherry by NheI/BsrGI digestion from EF1a-mCherry-P2A-Hygro plasmid (Addgene #135003). A previously validated hiPSC line^46^ was co-transfected with 1 μg lentiCRISPR-v2 plasmid (Addgene, 52961) containing TPRX1 targeting sgRNA and 2 μg TPRX1-2A-mCherry-PGK-PuroR donor plasmid using lipofectamine 3000 according to manufacturer’s instructions. The next day, cells were subjected to 1 μg/mL puromycin for 5 days and seeded as single cells onto 96-well plates. Individual colonies were expanded and screened by genomic PCR. To generate the GATA3-mKO2 line, cells were transfect with pGG195/GATA3mKO2 and px459/GATA3 which were kindly gifted by Ge Guo^47^.

### Chemical Screens for 8CLC induction

Naïve TPRX1–mCherry PSCs were seeded at 20,000 cells per well on Geltrex-coated 48-well plates in HENSM supplemented with 10 µM Y-27632. Twenty-four hours after seeding, the medium was replaced with fresh medium without Y-27632. On day 3, cells were treated with individual compounds at the indicated concentrations for 48 h. Vehicle-treated cells were used as negative controls. Trichostatin A (TSA) and DZNep were included as positive controls based on previous reports demonstrating induction of ZGA-associated transcriptional programs^7^. Following treatment, cells were harvested with 5 min dissociation with Trypsin, resuspended in 3% FBS containing 0.5 mM EDTA/PBS and analyzed by flow cytometry. The proportion of mCherry-positive cells was quantified for each condition and normalized relative to DMSO controls.

### Rapid induction of 8CLC

Naive PSCs were maintained on Geltrex in the corresponding naïve culture medium and passaged according to standard protocols. Upon reaching approximately 60% confluency, the culture medium was replaced with PRBJN induction which consists of PY60 (7.5 μM), Retinoic acid (0.5 μM), Bomedemstat (1 μM), JQKD82 (1 μM), and Nutlin-3a (0.5 μM). Because Nutlin-3a induces p53-dependent cytotoxicity^48^, we titrated its concentration and identified 0.5 μM as a dose that preserved 8CLC-inducing activity without significantly reducing viability within two days (Fig. S2C). Medium was refreshed daily throughout the induction period. To improve cell viability during induction, caspase inhibitors Emricasan (5 μM) or QVD-OPh (20 μM) were included in the culture medium.

### Flow cytometry and sorting

For flow cytometry experiments, cells were dissociated into single cell suspensions with 5 min trypsin incubation and resuspended in %3 FBS containing 0.5 mM EDTA/PBS. Cells were analyzed in Cytoflex S (Beckman Coulter) flow cytometer. Data were analyzed using FlowJo. For cell sorting experiments, cells were dissociated into single cell suspensions with 5 min trypsin incubation. Pellets were resuspended in 3% FBS and 10 μM Y-27632 containing 0.5 mM EDTA/PBS, filtered using 40 μm strainers and sorted on a BD FACSAria III instrument.

### RNA isolation and quantitative RT–PCR

Total RNA was isolated using the NucleoSpin RNA kit (Macherey-Nagel, Germany) according to the manufacturer’s instructions. cDNA was synthesized using the iScript cDNA Synthesis Kit (Bio-Rad, USA), and quantitative PCR was performed using SyberGreen mix (Takara Bio, Japan) on a LightCycler 480 real-time PCR system (Roche, Switzerland). Primer sequences are provided in Supplementary Table 2.

### Bulk RNA sequencing and analysis

TPRX1–mCherry-positive cells were isolated by fluorescence-activated cell sorting using a BD FACSAria III instrument. Total RNA was extracted using the NucleoSpin RNA isolation kit (Macherey-Nagel, Germany). Libraries were generated using the TruSeq Stranded mRNA Library Preparation Kit (Illumina, USA) and sequenced on an Illumina NovaSeq 6000 platform. Raw sequencing reads were trimmed using Trimmomatic and aligned to the human reference genome (GRCh38, GENCODE v48 annotation) using Salmon in quasi-mapping mode. Transcript-level quantifications were imported into R using the tximport package and summarized to gene level. Differential expression analysis was performed using DESeq2.

### Single-cell RNA sequencing

Single-cell transcriptomic profiling was performed using the Parse Biosciences Evercode Whole transcriptome v3 kit. Cells from DMSO-treated and PRBJN-treated conditions were processed according to the manufacturer’s protocol. Raw sequencing data were processed using the Parse Biosciences pipeline (splite-pipe v1.6.1) and imported into Seurat (v4.3.0.1). Samples were processed individually and then merged using the Seurat merge() function. Cells containing fewer than 700 UMIs, fewer than 200 detected genes or more than 10% mitochondrial transcripts were excluded. Doublets were identified and removed using DoubletFinder (v2.0.3). Following quality control, data were log normalized using the NormalizeData function and the top 2,000 highly variable genes were identified using FindVariableFeatures. Data were scaled and principal component analysis (PCA) was performed. Uniform Manifold Approximation and Projection (UMAP) was computed using the first 30 principal components. Unsupervised clustering was performed using the Louvain algorithm at resolution 0.8 via FindNeighbors and FindClusters. Cluster identity was assigned based on the expression of known marker genes.

### Integration with human embryo single-cell datasets

Published single-cell RNA-seq datasets from human preimplantation embryos were integrated to generate a reference developmental landscape. Raw count matrices from Yan et al. (2013), Petropoulos et al. (2016) and Mazid et al. (2022) were processed individually in Seurat (v5.5.0). Each dataset was log normalised using NormalizeData followed by identification of the top 3,000 highly variable genes with FindVariableFeatures. Datasets were scaled and dimensionality reduction was performed using PCA. The three reference datasets were integrated using Seurat anchor-based integration to create a unified reference atlas^32,32,48,33,49^. UMAP coordinates were computed on the integrated embedding using RunUMAP to enable subsequent query projection. To project PRBJN-treated cells onto the reference atlas, DMSO or PRBJN cells were processed as a query dataset. Transfer anchors between the reference and the query were identified using FindTransferAnchors. Query cells were projected onto the reference UMAP using MapQuery, with developmental stage labels transferred from the reference atlas. The proportion of DMSO-and PRBJN-treated cells mapping to each developmental stage was quantified and used to assess transcriptional similarity to human preimplantation embryo stages. Annotation of DMSO and PRBJN-treated cells was performed using the Seurat (v4.3.0.1). Each reference dataset was individually normalized, scaled, and processed for variable feature selection and Principal Component Analysis (PCA). Transfer anchors between each reference and the query dataset were identified using FindTransferAnchors across the first 30 principal components. Reference metadata annotations were then projected onto the query data using TransferData (dims = 1:30, k.weight = 10).

### Gene co-expression network analysis

Gene regulatory network (GRN) inference was performed using pySCENIC^33^. Co-expression modules were inferred from the normalized single-cell expression matrix using GRNBoost2. Transcription factor binding motif enrichment was assessed against the cisTarget database to identify regulons. Regulon activity was scored across individual cells using AUCell. Networks were visualized using custom R scripts with igraph and ggraph.

### CRISPR-mediated ANXA2 and YAP1 knockout

Guide RNAs targeting ANXA2 or YAP1 were designed and cloned into pLentiCRISPRv2-Puro (Addgene #52981). The following guide RNA sequences were used: for ANXA2, 5’-AACTGATTGACCAAGATGCT-3’ and 5’-ATACTAACTTTGATGCTGAG-3’; for YAP1, 5’-GCATCAGATCGTGCACGTCCG-3’, 5’-GATCAGACAACAACA-3’ and 5’-GACGTTCATCTGGGACAGCA-3’. Lentiviruses were produced in HEK293Ts and transduced into naïve pluripotent stem cells. Transduced cells were selected with puromycin, and protein depletion was confirmed by western blotting.

### Western blotting

Cells were lysed in RIPA buffer supplemented with protease inhibitor cocktail. Equal amounts of protein were separated by SDS–PAGE and transferred onto PVDF membranes. Membranes were blocked with 5% non-fat milk in PBST for 1 h at room temperature and incubated overnight at 4°C with primary antibodies against ANXA2 (1:2,000) and YAP1 (1:1,000). After washing with PBST, membranes were incubated with the appropriate HRP-conjugated secondary antibodies. Protein bands were detected using enhanced chemiluminescence (ECL) reagents according to the manufacturer’s instructions.

### ATAC-seq library preparation and data processing

ATAC-seq libraries were generated from DMSO-and PY60-treated naïve pluripotent stem cells using the ATAC-seq kit(Active Motif, cat# 53150) according to the manufacturer’s instructions. Sequencing reads were aligned to the GRCh38 reference genome using Bowtie2. Duplicate reads and reads mapping to mitochondrial DNA were removed prior to downstream analyses. To correct for Tn5 insertion bias, reads were shifted +4 bp on the forward strand and−5 bp on the reverse strand. Blacklisted genomic regions were excluded using the ENCODE hg38 blacklist.

Peak calling was performed using MACS2 with parameters optimized for ATAC-seq data. A consensus peak set was generated by merging peaks across all samples and read counts per consensus peak were quantified using featureCounts. To account for potential global shifts in chromatin accessibility, library size normalization was performed using a stable peak-based approach. Briefly, peaks showing consistent accessibility across all samples (stable peaks) were identified and used to derive normalization scale factors, which were then applied in DESeq2 to correct for library size differences. Regions with an adjusted p-value < 0.05 and absolute log₂ fold change > 1 were classified as gaining or losing accessibility. Normalized signal tracks were generated as bigWig files and visualized using the Integrative Genomics Viewer (IGV).

### Motif enrichment analysis

Transcription factor motif enrichment analysis was performed on gained and lost peak sets using AME from the MEME Suite, with JASPAR2024 CORE vertebrate motifs as the reference database. All consensus peak sequences were used as the background model. A second-order Markov background model was estimated from background sequences using fasta-get-markov. To identify transcription factor binding motifs enriched at promoters of differentially expressed genes, promoter sequences were extracted for genes upregulated more than 2-fold following PY60 treatment or DUX4 overexpression relative to DMSO controls. Promoter regions were defined as ±2 kb windows centered on the transcription start site of each gene. Genomic coordinates of promoter regions were retrieved based on Ensembl GRCh38 gene annotations, and corresponding DNA sequences were extracted from the hg38 reference genome using bedtools getfasta. Motif enrichment analysis was performed using AME from the MEME Suite with JASPAR2024 CORE vertebrate motifs as the reference database. Promoter sequences of all expressed genes were used as the background sequence set, and a second-order Markov background model was estimated using fasta-get-markov. Enriched motifs were ranked by q-value, and the percentage of target versus background sequences containing each motif was calculated to assess the specificity of enrichment.

### Immunofluorescence staining

Cells were fixed in 4% paraformaldehyde for 15 min at room temperature (RT) and washed three times with PBS. Cells were then permeabilized with 0.5% Triton X-100 in PBS for 10 min and blocked in PBS containing 3% serum for 1 hr at RT. Primary antibodies anti-TPRX1 (Sigma, WH0284355M2) 1:500, Anti-OCT4 (Abcam, ab19857) 1:250, Anti-KLF17 (Sigma, HPA024629) 1:250, LEUTX (Novus Biologicals, NBP1-90890) 1ug/ml were incubated overnight at 4°C followed by fluorophore-conjugated secondary antibodies (Goat anti-Mouse IgG Alexa Fluor 488 (Thermo Fisher, cat# A-11001) or Goat anti-Rabbit IgG Alexa Fluor 488 (Thermo Fisher, cat# A11034). Nuclei were counterstained with 1 μg/ml DAPI for 5 min. Images were acquired using LEICA SP8 confocal microscope and processed using ImageJ. Blastoids were transferred to BSA-coated plates containing 4% paraformaldehyde (PFA) and fixed for 15 min at room temperature. Following fixation, blastoids were blocked and permeabilized in PBS containing 3% BSA and 0.05% Triton X-100 for 4 h followed by transfer onto 40 μL droplets containing primary antibodies against NANOG (1:200) and incubated at 37°C for 2 h. To prevent evaporation, antibody droplets were surrounded with a hydrophobic barrier pen and covered with PBS. After primary antibody incubation, blastoids were washed three times by sequential transfer into 100 μL PBS droplets and subsequently incubated in 30 μL droplets containing the appropriate fluorophore-conjugated secondary antibodies for 1 h at room temperature. Following three additional PBS washes, blastoids were imaged using a Leica SP8 confocal microscope.

### Trophoblast stem cell differentiation assay

GATA3-mKO2 reporter cells were treated with either DMSO or PRBJN for 24 h and transferred to N2B27 basal medium and cultured for an additional four days without continued compound exposure. Differentiation into STBs was performed using both suspension and adherent culture conditions. For suspension differentiation, TSCs were seeded at 50,000 cells per well in AggreWell 400 plates (StemCell Technologies) using STB medium (Advanced DMEM/F12 supplemented with GlutaMAX, 0.1 mM 2-mercaptoethanol, penicillin– streptomycin, N2, B27, 50 ng ml⁻¹ human EGF, 2 µM CHIR99021, 0.8 mM valproic acid, and 5 µM Y27632). In parallel, TSCs were seeded onto Geltrex-coated plates and differentiated using the same STB medium under adherent conditions. Medium was refreshed every day. On day 5, conditioned media were collected and analyzed for hCG secretion using a commercially available qualitative hCG pregnancy test according to the manufacturer’s instructions. For EVT differentiation, TSCs were plated at 300,000 cells per well on Matrigel-coated 12-well plates using EVT medium (Advanced DMEM/F12 supplemented with GlutaMAX, 0.1 mM 2-mercaptoethanol, penicillin–streptomycin, N2, B27, and 5 µM Y27632). Medium was refreshed daily. Cells were collected on day 5 for downstream analyses.

### Induction of Blastoids

Blastoid generation experiments was approved by the Koç University Biomedical Research Ethics Committee (2025.068.IRB2.031) and followed The International Society for Stem Cell Research (ISSCR) Guidelines for Stem Cell Research and Clinical Translation^50^. Naive pluripotent stem cells (PSCs) were maintained in HENSM medium on Geltrex-coated plates. Following treatment with either 0.1% DMSO or PRBJN for 12 h, cells were dissociated using 1 mM EDTA in PBS for 10–15 min at room temperature. AggreWell 400 plates were prepared by treatment with Anti-Adherence Rinsing Solution, followed by centrifugation at 500 × g for 1 min, sonication for 2 min to remove trapped air bubbles, and incubation at room temperature for at least 1 h. Wells were then washed twice with N2B27 medium. Cells were collected by centrifugation and resuspended in aggregation medium consisting of N2B27 supplemented with 0.3% BSA and 1× CEPT cocktail and seeded at a density of 45 cells per microwell. Plates were incubated at room temperature for 15 min and centrifuged at 300 × g for 3 min. Aggregates were cultured in aggregation medium supplemented with 1× CEPT cocktail for the first 24 hours after which, half of the medium was replaced with 2× PALLY medium consisting of N2B27 supplemented with PD0325901 (2 μM), A83-01 (2 μM), lysophosphatidic acid (LPA; 5 μM), human LIF (80 ng/mL), and 1× CEPT cocktail. On day 2, the medium was completely replaced with 1× PALLY medium. On days 3 and 4, cultures were maintained in N2B27 supplemented with LPA, GA-017 (2.5 μM), and 1× CEPT cocktail. Thereafter, the medium was refreshed every 24 h with the same formulation for the remainder of the culture period. All cultures were maintained under hypoxic conditions (5% O₂ and 5% CO₂) throughout blastoid generation.

### Mouse embryo injection

Mouse embryo injection experiments were performed under a protocol approved by Koç University Animal Ethics Committee and in accordance with institutional animal care and use guidelines. Naïve PSCs stably expressing H2B–mCherry under a constitutive promoter were used as donor cells to enable tracking of injected human cells within host embryos. Donor cells were treated with either 0.1% DMSO or the PRBJN for 12 hours, after which cultures were dissociated to single-cell suspensions using 0.05% trypsin for 5 mins at 37°C and resuspended in PBS supplemented with 3% FBS. CB6F1 mice were used for embryo generation. Embryos were collected at embryonic day 0.5 (E0.5) and cultured in Global® (LG) drops covered with mineral oil under standard conditions (37°C, 5% CO₂) until they reached the 8-cell stage at approximately E2.5. Approximately 8–10 donor human cells were microinjected beneath the zona pellucida per embryo. Following injection, embryos were returned to Global® medium and cultured for an additional 24 hours until blastocyst formation. Blastocysts were imaged using fluorescence microscopy.

## Data availability

All sequencing data generated in this study have been deposited in the Gene Expression Omnibus (GEO) under the accession numbers GSE335948 (RNA-seq), GSE338528 (ATAC-seq), and GSE338782 (scRNA-seq). Publicly available human embryo RNA-seq datasets were obtained from GEO (GSE36552) and ArrayExpress (E-MTAB-3929). Publicly available 8CLC datasets were obtained from the CNGB Nucleotide Sequence Archive under accession number CNP0001454.

## Code availability

The computational analyses were performed using an in-house analysis pipeline, which is publicly available at https://github.com/onderlab/Chemical_induction_8CLC.

## Acknowledgements

We thank Jun Qi (Dana Farber Cancer Institute) for providing JQKD82, Derya Deniz Özdemir for assistance with mouse embryo injections and Serçin Karahüseyinoğlu for help with blastoid imaging. The authors gratefully acknowledge the use of the services and facilities of the Koç University Research Center for Translational Medicine (KUTTAM).

## Author contributions

A.O. and T.T.O. conceived ideas; A.O. generated TRPX1-reporter lines, performed chemical screening, validation, bulk RNA and ATAC-sequencing and associated bioinformatic analyses; E.O. generated *TP53* knockout cell lines and contributed to experimental validation; N.K. generated GATA3-mKO2 reporter cell line; S.U. performed differentiation assays and generated blastoids; K.H., M.L and C.A performed single cell sequencing and bioinformatics analyses; A.O and T.T.O. wrote the manuscript.

## Materials & Correspondence

Correspondence to Tamer T. Onder

---

**Figure S1.**
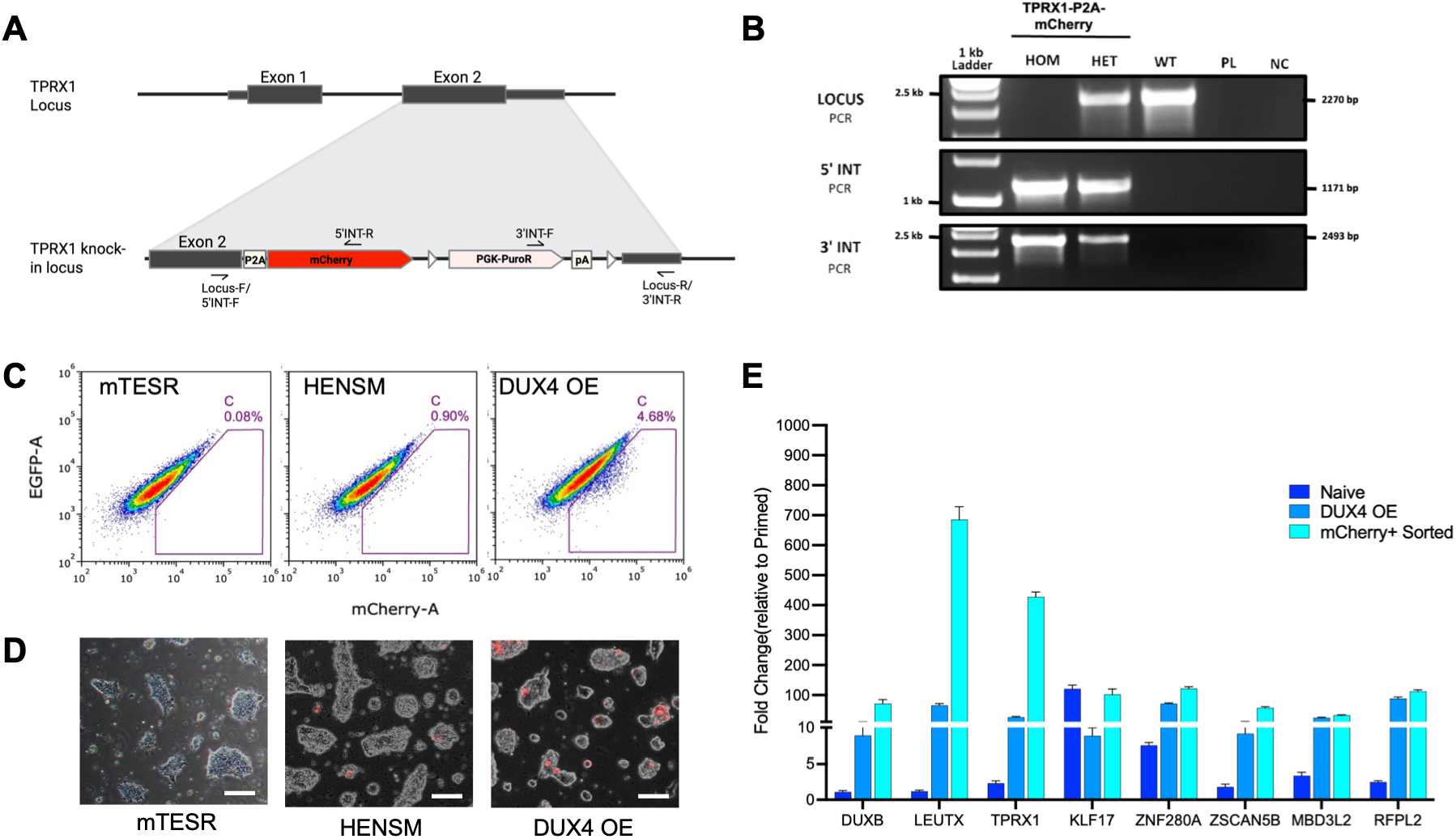
**A**, Schematic of the TPRX1 locus and the knock-in strategy. Primer positions used for genotyping (Locus F/R, 5’INT F/R and 3’INT F/R) are indicated. **B,** Genotyping PCR confirming successful knock-in of the TPRX1–P2A–mCherry cassette. Locus PCR, 5’ integration PCR and 3’ integration PCR are shown for homozygous (HOM), heterozygous (HET) and wild-type (WT) clones, alongside a plasmid control (PL) and no-template negative control (NC). **C,** Representative flow cytometry plots showing the proportion of mCherry-positive cells under primed (mTeSR1), naïve (HENSM) and DUX4-overexpressing conditions. **D,** Representative fluorescence microscopy images of TPRX1–mCherry reporter cells under mTeSR1, HENSM and DUX4-overexpressing conditions. Scale bars, 100 um. **E,** RT–qPCR analysis of established 8CLC-associated marker genes in naïve, DUX4-overexpressing and mCherry-positive sorted cells

**Figure S2.**
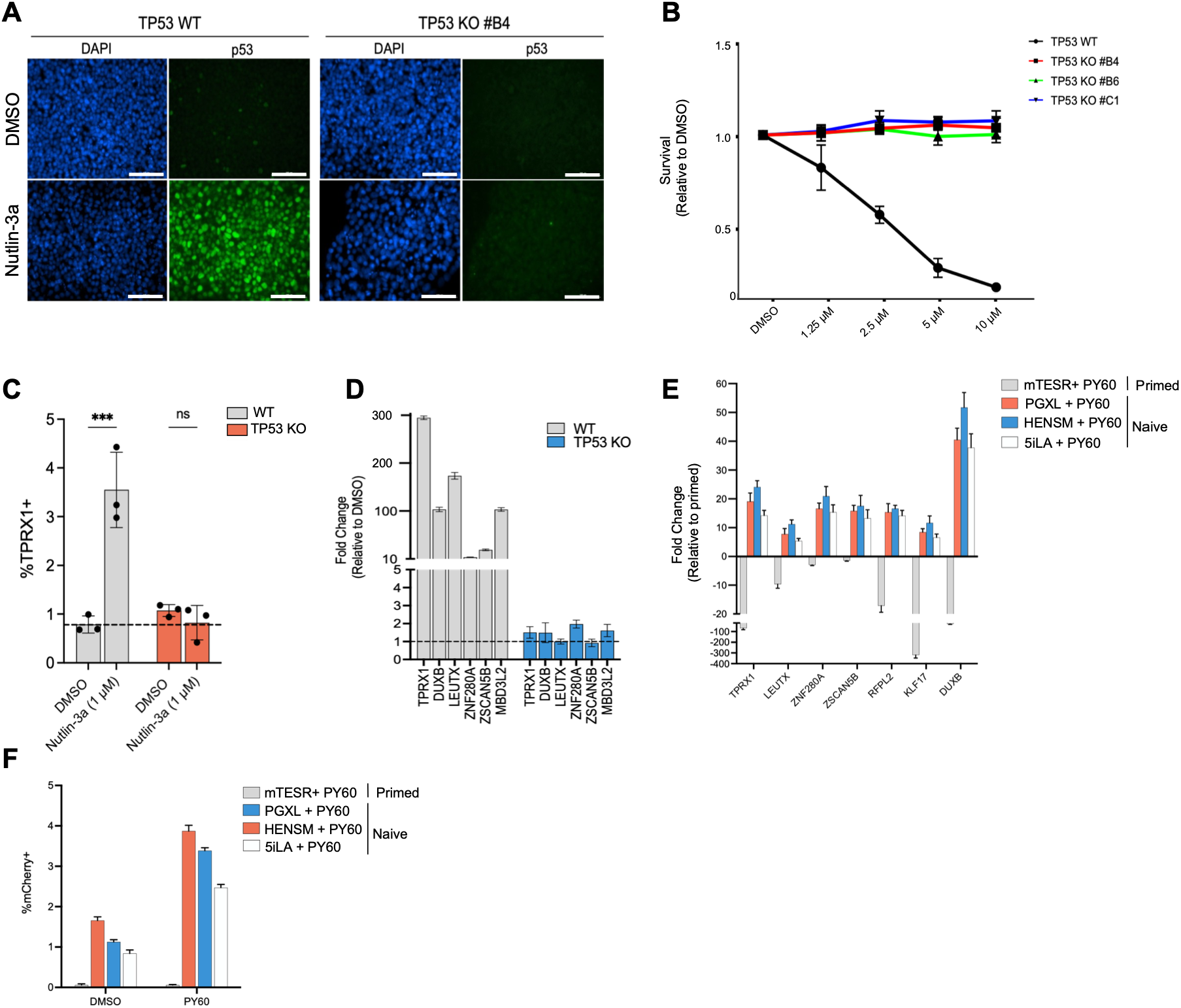
**A**, Immunofluorescence staining of p53 (green) and DAPI (blue) in TP53 WT and TP53 KO #B4 cells treated with DMSO or Nutlin-3a (5 µM). Scale bar, 100 µm. **B,** Cell viability of WT and TP53 KO clones treated with increasing concentrations of Nutlin-3a (1.25, 2.5, 5, 10) for 2 days, measured relative to DMSO-treated controls. **C,** Quantification of the proportion of TPRX1–mCherry-positive cells in TP53 WT and TP53 KO clones following treatment with DMSO and Nutlin-3a(1µM). Data represent mean ± SD (n=3). ***p < 0.001; ns, not significant. **D,** RT–qPCR analysis of representative 8CLC-associated marker genes following treatment with Nutlin-3a in TP53 WT and TP53 KO #B4 cells.. **E,** RT–qPCR analysis of 8CLC-marker genes following PY60 treatment across multiple naïve and primed culture conditions F, Quantification of TPRX1-positive cells under the culture conditions shown in panel E.

**Figure S3.**
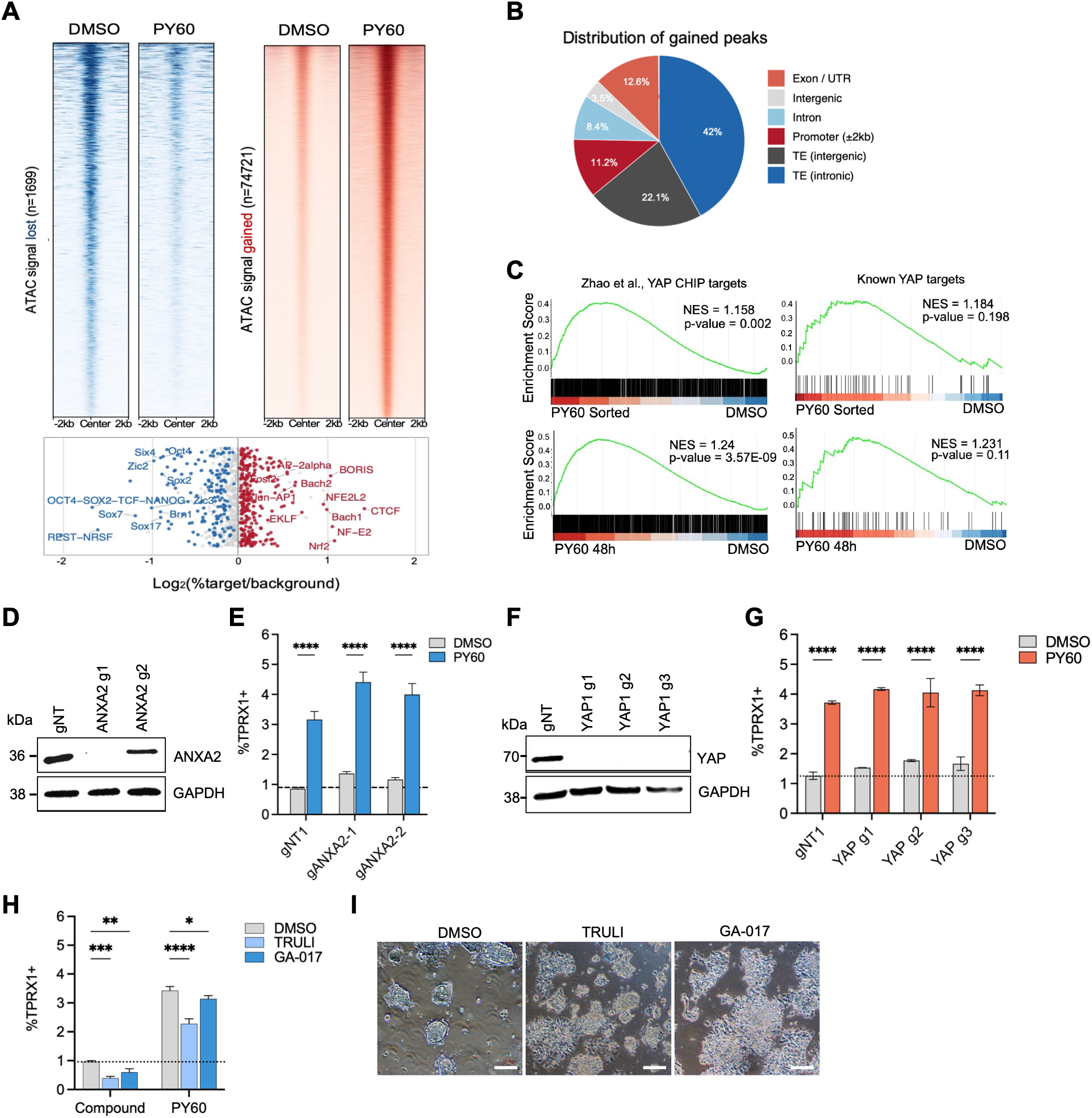
**A**, ATAC-seq heatmaps showing lost (n=1,699) and gained (n=74,721) chromatin accessibility peaks in DMSO and PY60-treated cells, centered at peak summits ±2 kb. Bottom, motif enrichment analysis (HOMER) showing transcription factor binding motifs enriched in lost (blue) or gained (red) chromatin regions, plotted as Log₂(%target/background). **B,** Pie chart showing the genomic distribution of gained ATAC-seq peaks across annotated regions including promoters (±2 kb), exons/UTRs, introns, intergenic regions, and transposable elements (TEs). **C,** Gene set enrichment analysis plots showing enrichment of YAP ChIP-seq target genes (Zhao et al., left) and known YAP target genes (right) in PY60-sorted mCherry-positive cells (top) and PY60-treated bulk cells at 48 hours (bottom) relative to DMSO controls. **D,** Immunoblots confirming ANXA2 depletion in cells transduced with two independent ANXA2-targeting guide RNAs (gANXA2-1 and gANXA2-2) compared with non-targeting control (gNT1). **E,** Quantification of the proportion of TPRX1-positive cells in ANXA2-depleted and control cells treated with DMSO or PY60. Statistical significance was assessed by two-way ANOVA. ****p < 0.0001. **F,** Immunoblots confirming YAP1 depletion in cells transduced with three independent YAP1-targeting guide RNAs (YAP1 g1, g2, g3) compared with non-targeting control (gNT1). **G,** Quantification of the proportion of TPRX1-positive cells in YAP1-depleted and control cells treated with DMSO or PY60. Statistical significance was assessed by two-way ANOVA. ****p < 0.0001. **H,** Quantification of the proportion of TPRX1-positive cells following treatment with the LATS1/2 inhibitors TRULI(10 μM) and GA-017(10 μM), alone or in combination with PY60. Statistical significance was assessed by two-way ANOVA. *p < 0.05, **p < 0.01, ***p < 0.001, ****p < 0.0001.. I, Representative bright-field images of naïve cells treated with DMSO, TRULI or GA-017.Scale bars, 200 μm.

**Figure S4.**
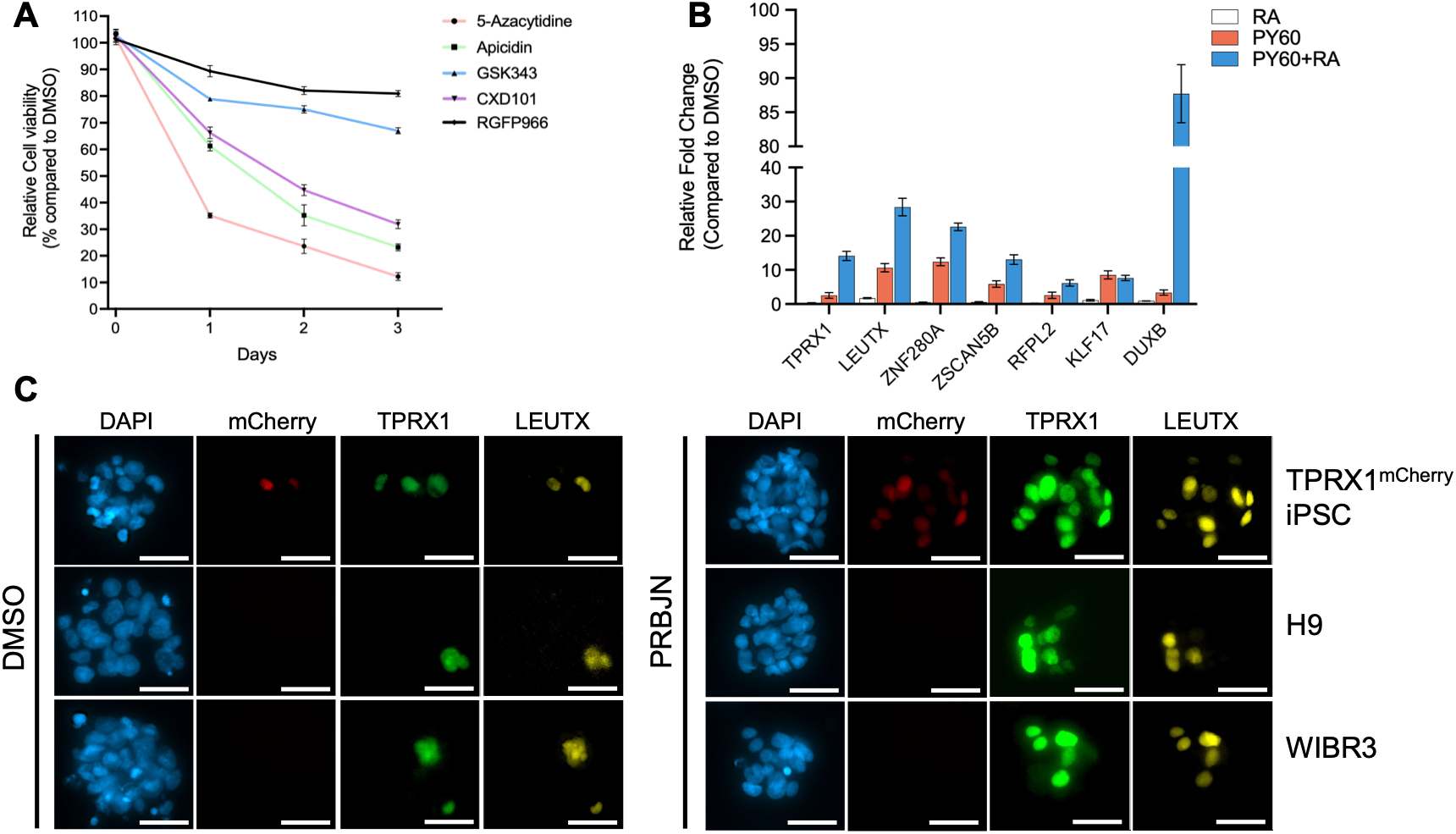
**A**, Cell viability analysis of naïve cells treated with selected secondary screen hit compounds over a three-day time course. **B,** RT– qPCR analysis of 8CLC-associated marker genes following treatment with 0.5 µM retinoic acid (RA), 5 µM PY60 or PY60 combined with RA. **C,** Immunofluorescence staining of PRBJN or DMSO-treated TPRX1-mCherry iPSC, H9 and WIBR3 lines for representative 8CLC-associated markers, LEUTX and TPRX1. Scale bars, 50 μm.

**Figure S5.**
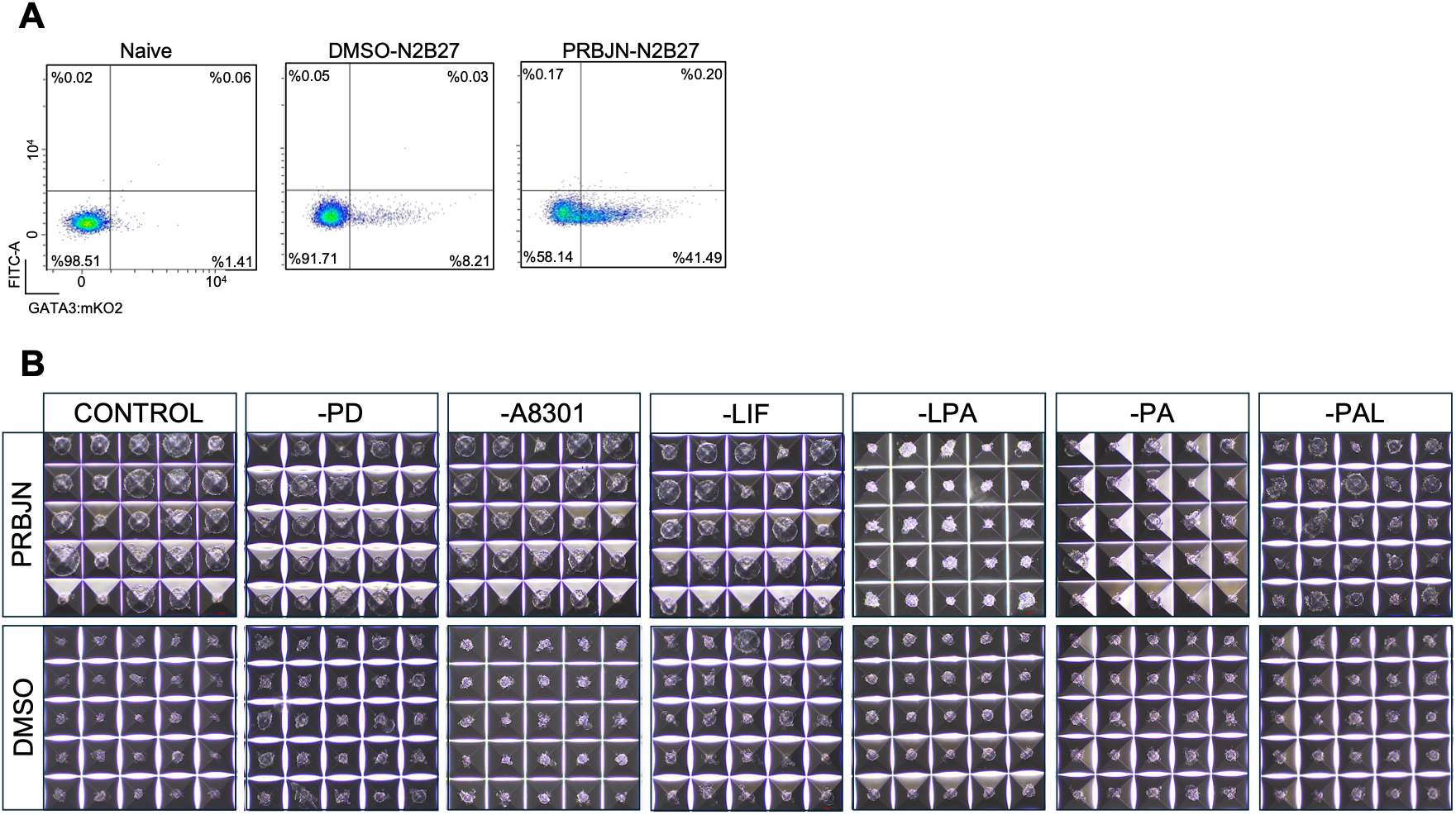
**A**, Representative flow cytometry plots showing the proportion of GATA3–mKO2-positive cells in untreated naïve cells (left), DMSO-treated cells following N2B27 differentiation (middle) and PRBJN-treated cells following N2B27 differentiation (right). **B,** Representative bright-field images of microwell arrays showing blastoid formation from naïve cells pre-treated with PRBJN (top) or DMSO (bottom) for 12 h, under the complete PALLY condition and individual component dropouts (−PD, −LIF, −A8301, −LPA, −PA, −PAL). Each grid shows one microwell array per condition.

## References

1. De Paepe, C., Krivega, M., Cauffman, G., Geens, M. & Van de Velde, H. Totipotency and lineage segregation in the human embryo. Mol. Hum. Reprod. 20, 599–618 (2014).

2. Braude, P., Bolton, V. & Moore, S. Human gene expression first occurs between the four-and eight-cell stages of preimplantation development. Nature 332, 459–461 (1988).

3. Vassena, R. et al. Waves of early transcriptional activation and pluripotency program initiation during human preimplantation development. Development 138, 3699–3709 (2011).

4. Lee, M. T., Bonneau, A. R. & Giraldez, A. J. Zygotic Genome Activation During the Maternal-to-Zygotic Transition. Annu. Rev. Cell Dev. Biol. 30, 581–613 (2014).

5. Taubenschmid-Stowers, J. et al. 8C-like cells capture the human zygotic genome activation program *in vitro*. Cell Stem Cell 29, 449–459.e6 (2022).

6. Yu, X. et al. Recapitulating early human development with 8C-like cells. Cell Rep. 39, 110994 (2022).

7. Mazid, M. A. et al. Rolling back human pluripotent stem cells to an eight-cell embryo-like stage. Nature 605, 315–324 (2022).

8. Yoshihara, M. et al. Transient DUX4 expression in human embryonic stem cells induces blastomere-like expression program that is marked by SLC34A2. Stem Cell Rep. 17, 1743– 1756 (2022).

9. Yu, X. et al. Recapitulating early human development with 8C-like cells. Cell Rep. 39, 110994 (2022).

10. Mazid, M. A. et al. Rolling back human pluripotent stem cells to an eight-cell embryo-like stage. Nature 605, 315–324 (2022).

11. Vallot, A. & Tachibana, K. The emergence of genome architecture and zygotic genome activation. Curr. Opin. Cell Biol. 64, 50–57 (2020).

12. Schulz, K. N. & Harrison, M. M. Mechanisms regulating zygotic genome activation. Nat. Rev. Genet. 20, 221–234 (2019).

13. Pessina, P. et al. Selective RNA sequestration in biomolecular condensates directs cell fate transitions. Nat. Biotechnol. 1–16 (2025) doi:10.1038/s41587-025-02853-z.

14. Grow, E. J. et al. p53 convergently activates Dux/DUX4 in embryonic stem cells and in facioscapulohumeral muscular dystrophy cell models. Nat. Genet. 53, 1207–1220 (2021).

15. Zhang, W. et al. Zscan4c activates endogenous retrovirus MERVL and cleavage embryo genes. Nucleic Acids Res. 47, 8485–8501 (2019).

16. Li, S. et al. Capturing totipotency in human cells through spliceosomal repression. Cell 187, 3284–3302.e23 (2024).

17. Shalhout, S. Z. et al. YAP-dependent proliferation by a small molecule targeting annexin A2. Nat. Chem. Biol. 17, 767–775 (2021).

18. Bayerl, J. et al. Principles of signaling pathway modulation for enhancing human naive pluripotency induction. Cell Stem Cell 28, 1549–1565.e12 (2021).

19. Bredenkamp, N. et al. Wnt Inhibition Facilitates RNA-Mediated Reprogramming of Human Somatic Cells to Naive Pluripotency. Stem Cell Rep. 13, 1083–1098 (2019).

20. Fischer, L. A., Khan, S. A. & Theunissen, T. W. Induction of Human Naïve Pluripotency Using 5i/L/A Medium. Methods Mol. Biol. 2416, 13–28 (2022).

21. Göke, J. et al. Dynamic transcription of distinct classes of endogenous retroviral elements marks specific populations of early human embryonic cells. Cell Stem Cell 16, 135–141 (2015).

22. Xiang, Y. et al. Endogenous retroviruses synthesize heterologous chimeric RNAs to reinforce human early embryo development. Science 391, eadv5257 (2026).

23. Zhao, B. et al. TEAD mediates YAP-dependent gene induction and growth control. Genes Dev. 22, 1962–1971 (2008).

24. Cordenonsi, M. et al. The Hippo transducer TAZ confers cancer stem cell-related traits on breast cancer cells. Cell 147, 759–772 (2011).

25. Viukov, S. et al. Human primed and naïve PSCs are both able to differentiate into trophoblast stem cells. Stem Cell Rep. 17, 2484–2500 (2022).

26. Dattani, A., Huang, T., Liddle, C., Smith, A. & Guo, G. Suppression of YAP safeguards human naïve pluripotency. Development 149, dev200988 (2022).

27. Yang, M. et al. Chemical-induced chromatin remodeling reprograms mouse ESCs to totipotent-like stem cells. Cell Stem Cell 29, 400–418.e13 (2022).

28. Xu, Y. et al. Derivation of totipotent-like stem cells with blastocyst-like structure forming potential. Cell Res. 32, 513–529 (2022).

29. Zhu, X. et al. N6-methyladenosine on L1PA governs the trans-silencing of LTRs and restrains totipotency in naive human embryonic stem cells. Cell Stem Cell 32, 1773–1791.e13 (2025).

30. Cossec, J.-C. et al. SUMO Safeguards Somatic and Pluripotent Cell Identities by Enforcing Distinct Chromatin States. Cell Stem Cell 23, 742–757.e8 (2018).

31. Yan, L. et al. Single-cell RNA-Seq profiling of human preimplantation embryos and embryonic stem cells. Nat. Struct. Mol. Biol. 20, 1131–1139 (2013).

32. Petropoulos, S. et al. Single-Cell RNA-Seq Reveals Lineage and X Chromosome Dynamics in Human Preimplantation Embryos. Cell 165, 1012–1026 (2016).

33. Aibar, S. et al. SCENIC: single-cell regulatory network inference and clustering. Nat. Methods 14, 1083–1086 (2017).

34. Guo, G. et al. Human naive epiblast cells possess unrestricted lineage potential. Cell Stem Cell 28, 1040–1056.e6 (2021).

35. Kagawa, H. et al. Human blastoids model blastocyst development and implantation. Nature 601, 600–605 (2022).

36. Iturbide, A. et al. Retinoic acid signaling is critical during the totipotency window in early mammalian development. Nat. Struct. Mol. Biol. 28, 521–532 (2021).

37. Macfarlan, T. S. et al. Embryonic stem cell potency fluctuates with endogenous retrovirus activity. Nature 487, 57–63 (2012).

38. Yang, F. et al. DUX-miR-344-ZMYM2-Mediated Activation of MERVL LTRs Induces a Totipotent 2C-like State. Cell Stem Cell 26, 234–250.e7 (2020).

39. Wang, Y. et al. LSD1 is a subunit of the NuRD complex and targets the metastasis programs in breast cancer. Cell 138, 660–672 (2009).

40. Ohguchi, H. et al. Lysine Demethylase 5A is Required for MYC Driven Transcription in Multiple Myeloma. Blood Cancer Discov. 2, 370–387 (2021).

41. Kong, X. et al. OTX2 inhibits human pluripotent stem cell reprogramming toward 8-cell-like and morula-like states. Nat. Commun. 17, 1685 (2026).

42. Su, X. et al. NELFA supports naïve pluripotency and drives 8C-like state in human embryonic stem cells. Nat. Commun. (2026) doi:10.1038/s41467-026-72714-z.

43. Bayerl, J. et al. Principles of signaling pathway modulation for enhancing human naive pluripotency induction. Cell Stem Cell 28, 1549–1565.e12 (2021).

44. Yanagida, A. et al. Naive stem cell blastocyst model captures human embryo lineage segregation. Cell Stem Cell 28, 1016–1022.e4 (2021).

45. Theunissen, T. W. et al. Systematic identification of culture conditions for induction and maintenance of naive human pluripotency. Cell Stem Cell 15, 471–487 (2014).

46. Alici-Garipcan, A. et al. NLRP7 plays a functional role in regulating BMP4 signaling during differentiation of patient-derived trophoblasts. Cell Death Dis. 11, 658 (2020).

47. Guo, G. et al. Human naive epiblast cells possess unrestricted lineage potential. Cell Stem Cell 28, 1040–1056.e6 (2021).

48. Vassilev, L. T. et al. In vivo activation of the p53 pathway by small-molecule antagonists of MDM2. Science 303, 844–848 (2004).

49. Stuart, T. et al. Comprehensive Integration of Single-Cell Data. Cell 177, 1888–1902.e21 (2019).

50. Lovell-Badge, R. et al. ISSCR Guidelines for Stem Cell Research and Clinical Translation: The 2021 update. Stem Cell Rep. 16, 1398–1408 (2021).

