## Supplemental Figures for "Rapid and efficient generation of human 8-cell-like cells for embryo modelling"

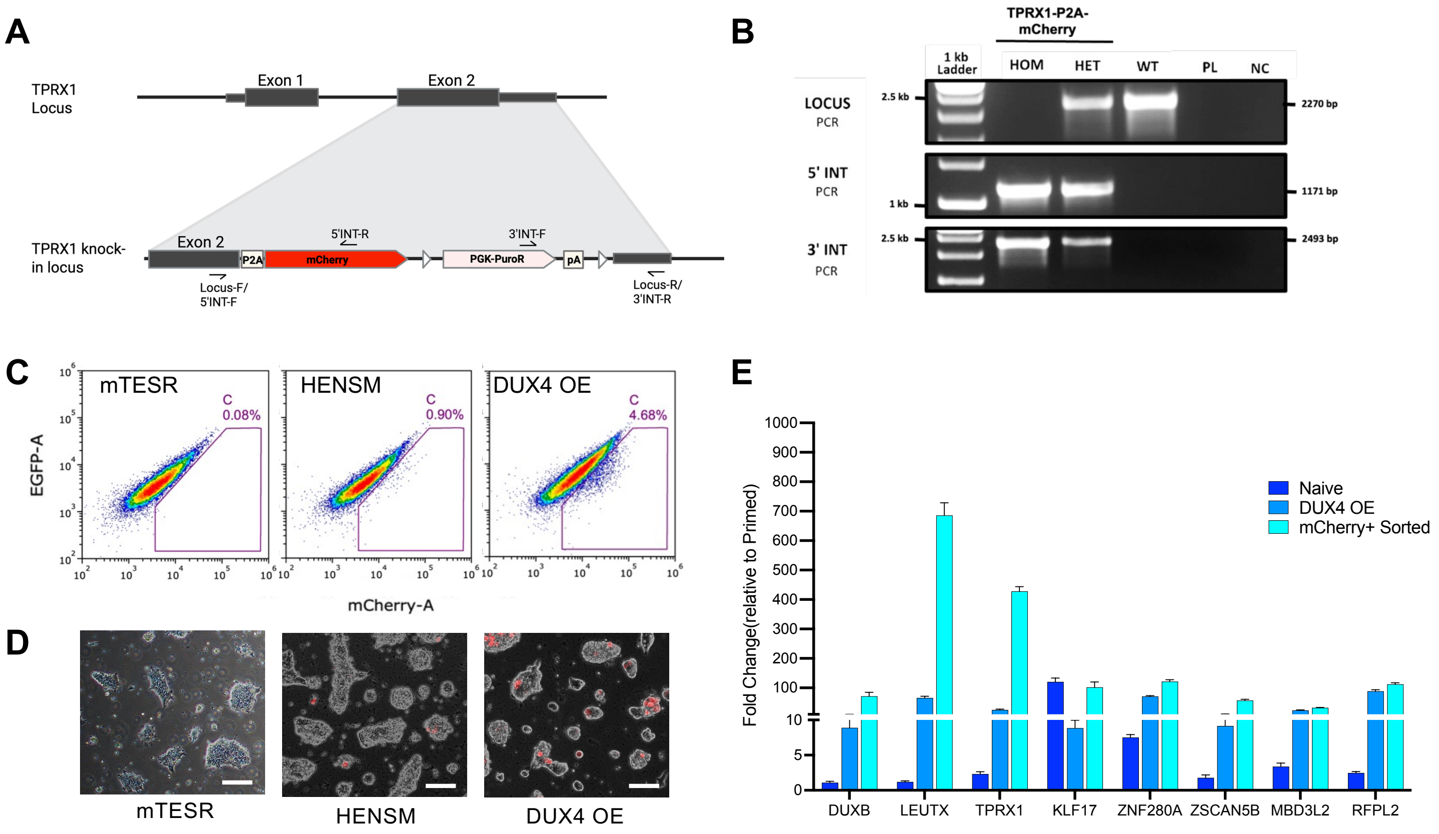


**Figure S1**

**A**, Schematic of the TPRX1 locus and the knock-in strategy. Primer positions used for genotyping (Locus F/R, 5'INT F/R and 3'INT F/R) are indicated. **B,** Genotyping PCR confirming successful knock-in of the TPRX1–P2A–mCherry cassette. Locus PCR, 5' integration PCR and 3' integration PCR are shown for homozygous (HOM), heterozygous (HET) and wild-type (WT) clones, alongside a plasmid control (PL) and no-template negative control (NC). **C**, Representative flow cytometry plots showing the proportion of mCherry-positive cells under primed (mTeSR1), naïve (HENSM) and DUX4-overexpressing conditions. **D,** Representative fluorescence microscopy images of TPRX1–mCherry reporter cells under mTeSR1, HENSM and DUX4-overexpressing conditions. Scale bars, 100 um. **E**, RT–qPCR analysis of established 8CLC-associated marker genes in naïve, DUX4-overexpressing and mCherry-positive sorted cells

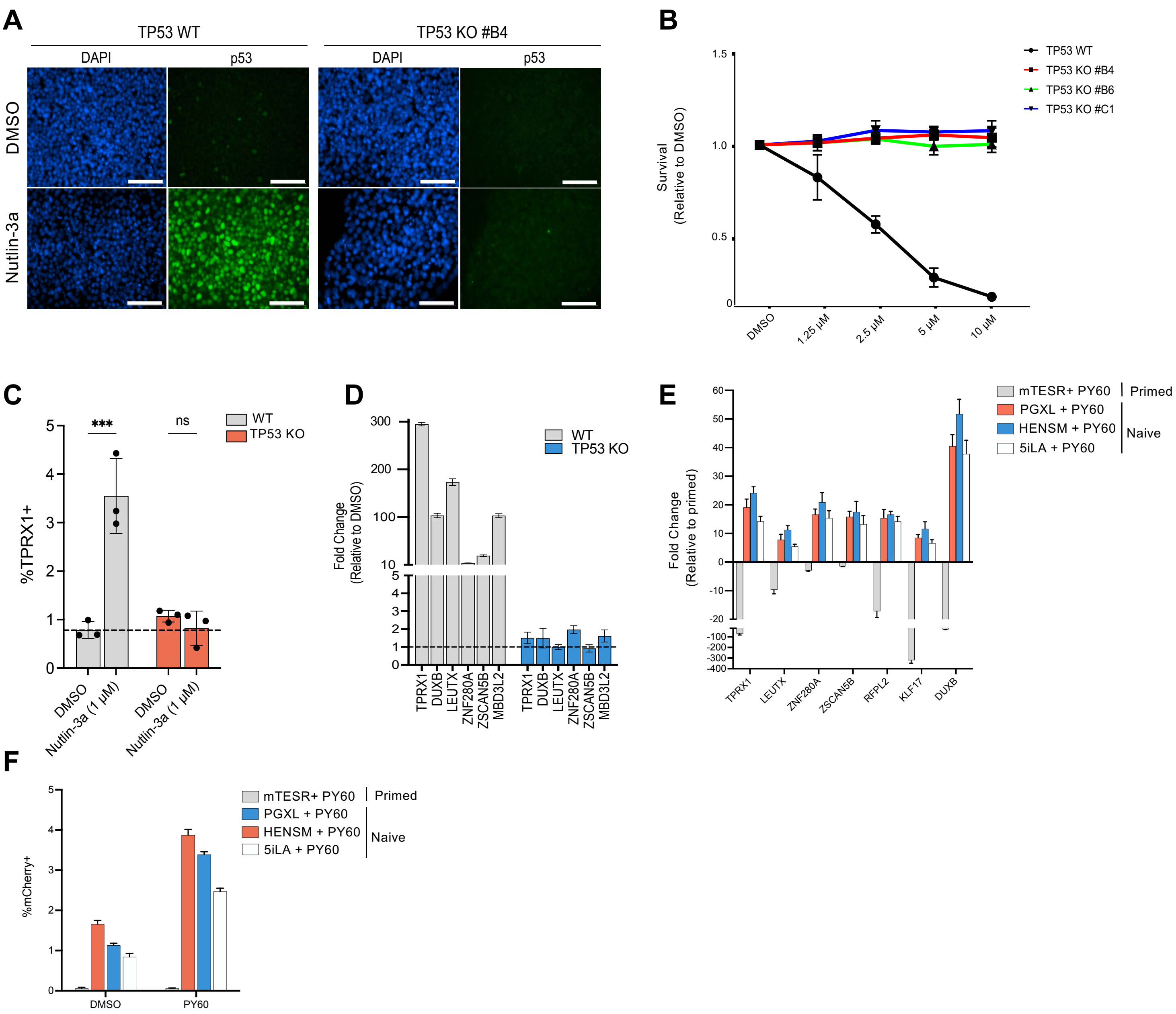


**Figure S2**

**A,** Immunofluorescence staining of p53 (green) and DAPI (blue) in TP53 WT and TP53 KO #B4 cells treated with DMSO or Nutlin-3a (5 µM). Scale bar, 100 µm. **B,** Cell viability of WT and TP53 KO clones treated with increasing concentrations of Nutlin-3a (1.25, 2.5, 5, 10) for 2 days, measured relative to DMSO-treated controls. **C,** Quantification of the proportion of TPRX1–mCherry-positive cells in TP53 WT and TP53 KO clones following treatment with DMSO and Nutlin-3a(1µM). Data represent mean ± SD (n=3). ***p < 0.001; ns, not significant. **D,** RT–qPCR analysis of representative 8CLC-associated marker genes following treatment with Nutlin-3a in TP53 WT and TP53 KO #B4 cells. **E,** RT–qPCR analysis of 8CLC-marker genes following PY60 treatment across multiple naïve and primed culture conditions **F,** Quantification of TPRX1-positive cells under the culture conditions shown in panel *E*.

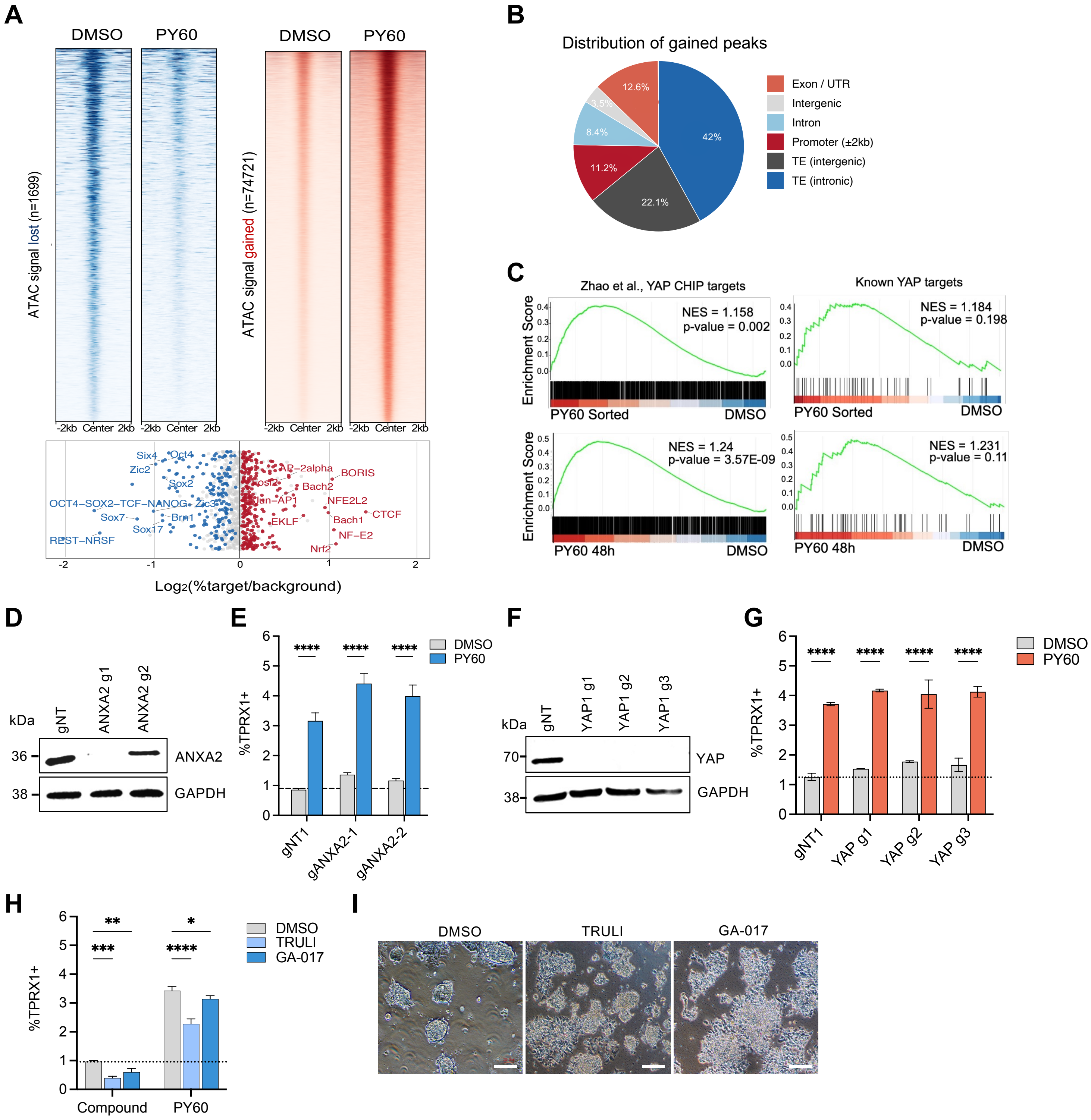


**Figure S3**

**A.** ATAC-seq heatmaps showing lost (n=1,699) and gained (n=74,721) chromatin accessibility peaks in DMSO and PY60-treated cells, centered at peak summits ±2 kb. Bottom, motif enrichment analysis (HOMER) showing transcription factor binding motifs enriched in lost (blue) or gained (red) chromatin regions, plotted as $Log_2$(%target/background). **B,** Pie chart showing the genomic distribution of gained ATAC-seq peaks across annotated regions including promoters (±2 kb), exons/UTRs, introns, intergenic regions, and transposable elements (TEs). **C,** Gene set enrichment analysis plots showing enrichment of YAP ChIP-seq target genes (Zhao et al., left) and known YAP target genes (right) in PY60-sorted mCherry-positive cells (top) and PY60-treated bulk cells at 48 hours (bottom) relative to DMSO controls. **D,** Immunoblots for ANXA2 depletion in cells transduced with two independent ANXA2-targeting guide RNAs (gANXA2-1 and gANXA2-2) compared with non-targeting control (gNT1). **E,** Proportion of TPRX1-positive cells in ANXA2-depleted and control cells treated with DMSO or PY60. Statistical significance was assessed by two-way ANOVA. ****$p < 0.0001$. **F,** Immunoblots for YAP1 depletion in cells transduced with three independent YAP1-targeting guide RNAs (YAP1 g1, g2, g3) compared with non-targeting control (gNT1). **G,** Proportion of TPRX1-positive cells in YAP1-depleted and control cells treated with DMSO or PY60. Statistical significance was assessed by two-way ANOVA. ****$p < 0.0001$. **H,** Proportion of TPRX1-positive cells following treatment with the LATS1/2 inhibitors TRULI(10 µM) and GA-017(10 µM), alone or in combination with PY60. Statistical significance was assessed by two-way ANOVA. *$p < 0.05$, **$p < 0.01$, ***$p < 0.001$, ****$p < 0.0001$.. **I,** Representative bright-field images of naïve cells treated with DMSO, TRULI or GA-017. Scale bars, 200 µm.

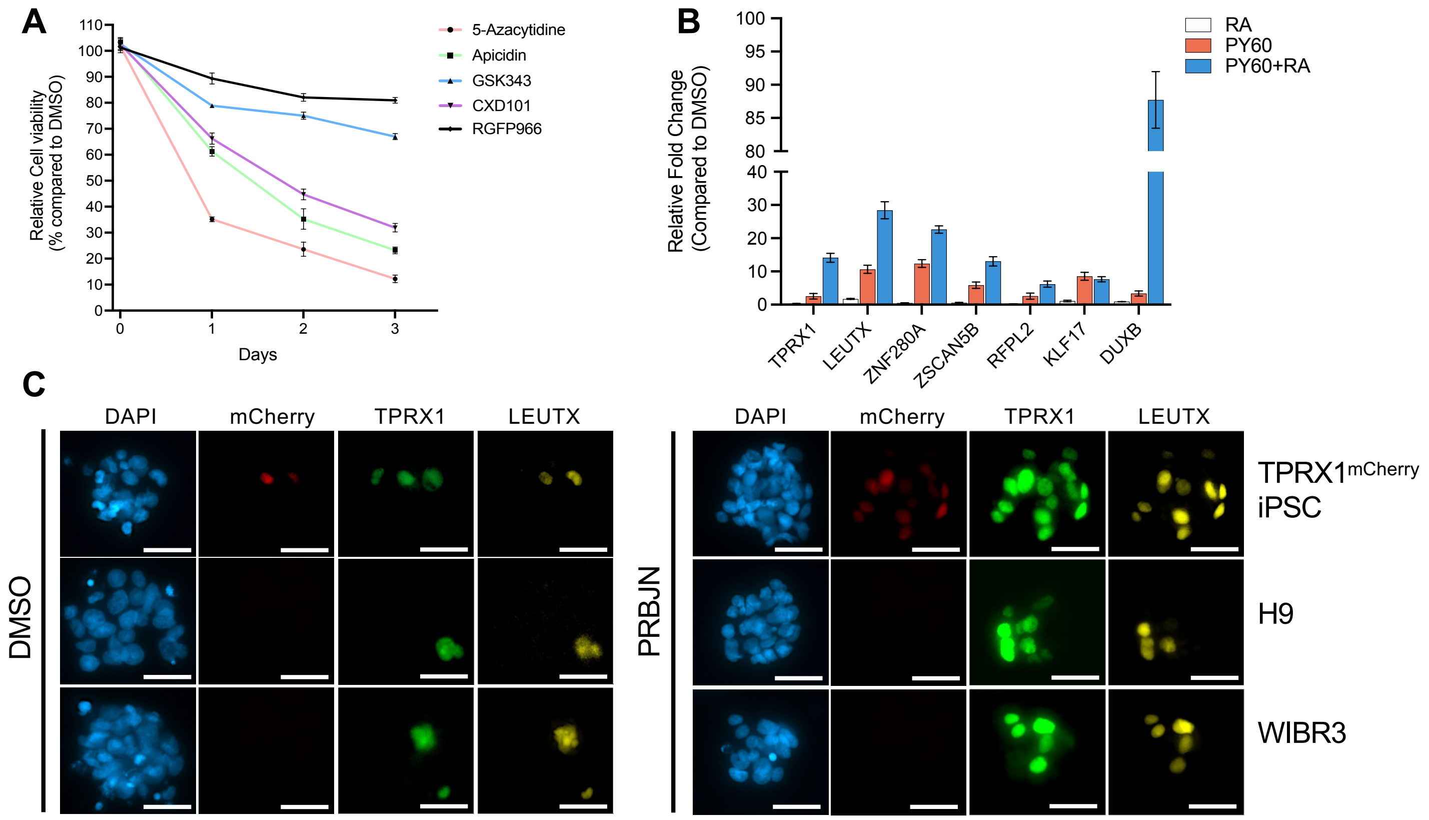


**Figure S4**

**A,** Cell viability analysis of naïve cells treated with selected secondary screen hit compounds over a three-day time course. **B,** RT–qPCR analysis of 8CLC-associated marker genes following treatment with 0.5 µM retinoic acid (RA), 5 µM PY60 or PY60 combined with RA. **C,** Immunofluorescence staining of PRBJN or DMSO-treated TPRX1-mCherry iPSC, H9 and WIBR3 lines for representative 8CLC-associated markers, LEUTX and TPRX1. Scale bars, 50 µm.

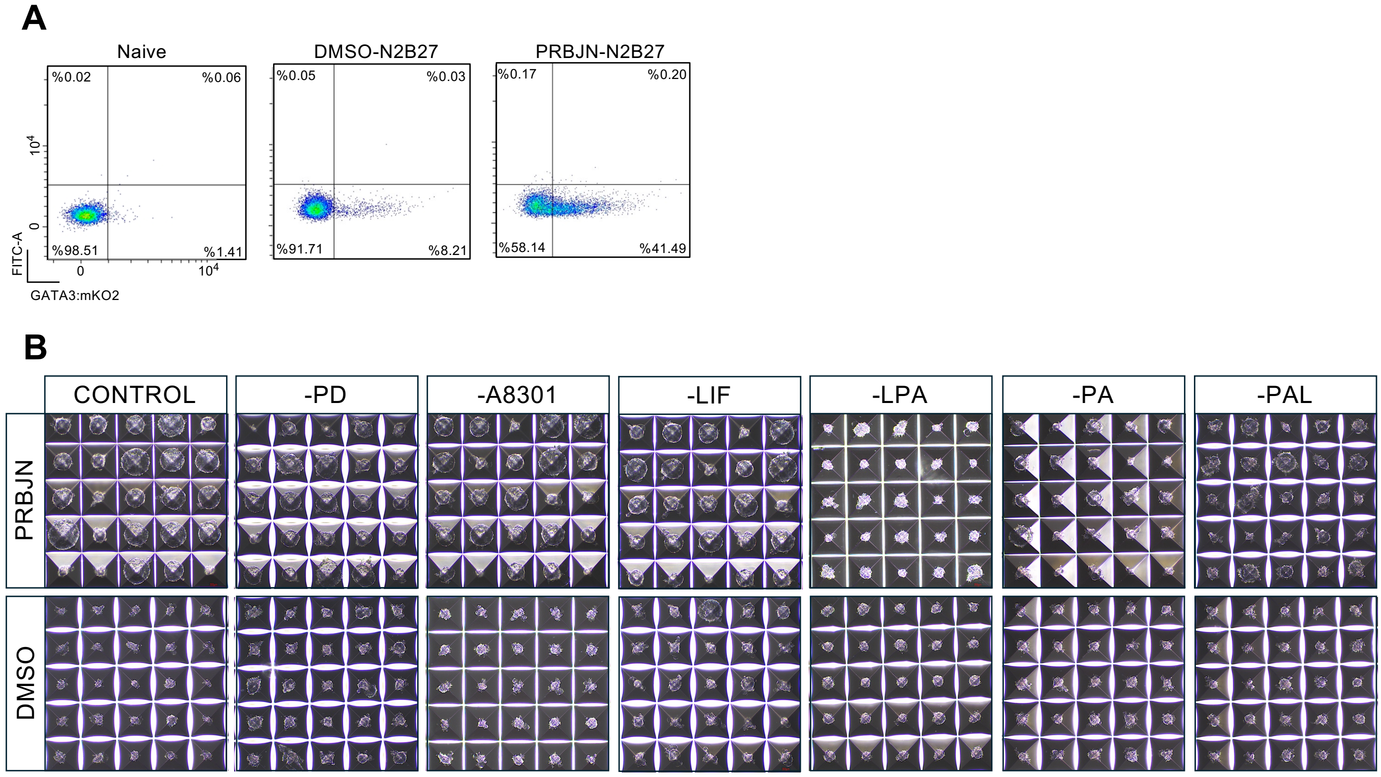


**Figure S5**

**A,** Representative flow cytometry plots showing the proportion of GATA3–mKO2-positive cells in untreated naïve cells (left), DMSO-treated cells following N2B27 differentiation (middle) and PRBJN-treated cells following N2B27 differentiation (right). **B,** Representative bright-field images of microwell arrays showing blastoid formation from naïve cells pre-treated with PRBJN (top) or DMSO (bottom) for 12 h, under the complete PALLY condition and individual component dropouts (−PD, −LIF, −A8301, −LPA, −PA, −PAL) on day 3. Each grid shows one microwell array per condition.
